# KDM3B enforces a non-mutational ErbB signaling state that sustains cholangiocarcinoma

**DOI:** 10.64898/2026.11.12.766134

**Authors:** Yuewan Luo, Orestis Nousias, Ekaterina Zhuravleva, Juan Lafuente-Barquero, Chirag Nepal, Antonio Marino, Supaporn Yangngam, Andrzej Taranta, Ana R. Colaço, Shufeng Lu, Michael Wierer, Colm J. O’Rourke, Jesper B. Andersen

## Abstract

More than 50% of cholangiocarcinoma (CCA) patients lack druggable mutations, showing broad non-mutational epigenetic perturbations, whose mechanistic insights and clinical benefit remain largely unexplored. Across multiple CCA patient cohorts, we identified prevalent significant upregulation of lysine demethylase (KDM) gene families, with KDM3B emerging as a critical epigenetic dependency for sustaining tumor fitness. *KDM3B* loss reduced colony formation, proliferation, and migration across intra- and extrahepatic CCA tumor models. Integrated multi-omics linked KDM3B to ErbB activation and downstream PI3K-AKT oncogenic signaling. Mechanistically, KDM3B is bound to the *ERBB3* promoter, removing repressive H3K9me2 marks to transcriptionally activate *ERBB3*, augmenting ErbB receptor activity. Spatial and single cell transcriptomics from patient tumors revealed that high KDM3B-ERBB3 coupling reshapes the tumor niche via enhancing the epithelial-mesenchymal transition crosstalk with neighborhood cells to establish a pro-invasive microenvironment. Pharmacologic inhibition *in vivo* of the KDM3B downstream ERBB-Akt regulatory axis suppressed CCA xenograft tumor growth. These findings establish KDM3B as a non-mutational epigenetic activator of ErbB-driven signaling, reshaping the tumor ecosystem, and defining the KDM3B-ErbB axis as a vulnerability in CCA.

**Highlights:**

- KDM3B establishes a non-mutational epigenetic dependency that sustains tumor fitness
- KDM3B activates ErbB signaling via H3K9me2-dependent chromatin regulation of *ERBB3*
- KDM3B-ERBB3 coupling reshapes the tumor ecosystem toward a pro-invasive state
- Targeting the KDM3B-ERBB-AKT axis suppresses CCA tumor growth

## Introduction

Cholangiocarcinoma (CCA) is an aggressive malignancy arising from the intra- and extrahepatic bile ducts, with 5-year overall survival less than 20%^1,2^. The incidence rates of CCA worldwide are expected to rise considerably regardless of significant molecular and clinical insights achieved in the past decade, including several approved therapies in the advanced tumor stages^2^. Still, this unmet clinical need warrants rethinking strategies, requiring biological understanding of how these tumors develop to enhance patient selection, advance therapies, and evade resistance mechanisms.

Non-mutational epigenetic reprogramming is increasingly recognized as a hallmark of cancer that establishes oncogenic transcriptional programs independently of DNA sequence alterations through coordinated epigenetic regulation at the chromatin, transcriptional, and post-transcriptional levels^3,4^. In CCA, beyond the well-characterized genetic alterations (*IDH1/2*, *FGFR2* fusions, *ERBB2* and *KRAS*) accounting for 30-40% of patients, for which targeted therapies are available today^2^, more than half of tumors are wildtype for these alterations, lacking key actionable drivers, but exhibiting pronounced enrichment and heterogeneity in epigenetic alterations^5,6^, with emerging evidence demonstrating their functional contribution to CCA tumor growth^7^.

Histone lysine demethylases (KDMs) represent a large class of chromatin modifiers, each responsible for controlling distinct histone marks and downstream transcriptional programs, which are often highlighted as deregulated in cancer^8^. KDMs span two mechanistic enzyme classes, including the FAD-dependent amine oxidases (KDM1/LSD) and Fe^2+^/2-oxoglutarate-dependent Jumonji C (JmjC) domain-containing protein (KDM2-7) subfamilies^8^. As proximal drivers, their biological roles make them therapeutically interesting; however, the highly conserved JmjC catalytic pocket limits selectivity and results in unwanted toxicities, suggesting their control of downstream regulatory pathways may discriminate clinical management^8,9^.

In CCA, recurrent mutations of KDMs are infrequent. However, several studies highlight augmented KDM expression and activity that promotes tumor growth^10–12^. In the liver, KDM3 shows context-dependency with elevated expression associated with PI3K-Akt signaling^13,14^. KDM3B, a key member of the KDM3 family, removes mono- and dimethyl marks from histone H3 lysine 9 (H3K9) and orchestrates transcriptional activation through interaction with the m6A reader (YTHDC1) to promote methylation^15,16^. Indeed, KDM3B was suggested to be predictive of poor recurrent free survival in liver cancers^13^. Despite these insights, the direct mechanistic role of KDM3B in non-mutational epigenetic reprogramming in tumors, specifically the underlying biology of its regulatory histone control, and therapeutic tractability remains largely unknown.

In multiple independent patient cohorts, we uncovered recurrent upregulation of *KDM3* and *KDM5* gene families in CCA tumors. We demonstrated that genetic *KDM3A/B* and *KDM5A/B* loss compromise tumor-associated phenotypes, with *KDM3B* showing dominance in tumor cell maintenance. Cellular and integrated molecular analyses revealed that KDM3B primarily exerts downstream oncogenic pathway control by activating ErbB signaling, directly regulating *ERBB3* receptor expression through H3K9me2 demethylation. Patient multi-omics further substantiated this regulatory axis, demonstrating that KDM3B-ERBB3 coupling reprograms tumor-stromal communication toward a pro-invasive niche. Pharmacological inhibition of ErbB signaling in xenograft models phenocopies *KDM3B* loss, achieving anti-tumor activity. Collectively, our study establishes KDM3B as a non-mutational epigenetic regulator that sustains CCA tumor cell maintenance by engaging ERBB3-mediated oncogenic ErbB signaling, reshaping a permissive tumor ecosystem.

## RESULTS

### Upregulation of KDM family members in cholangiocarcinoma

To delineate the contribution of epigenetic factors in cholangiocarcinoma (CCA), we analyzed the expression of 567 epigenetic genes^17^ in patient transcriptomes of 210 tumors versus 152 matched adjacent livers^18^, identifying in total 263 epigenetic factors significantly dysregulated in tumor tissues, of which a subset reduced cell viability in DepMap CRISPR screens (Figure 1A and S1A; Table S1A). Leveraging internal high-throughput drug screening data^5^, we evaluated the sensitivity of 8 CCA cell models to 61 selected epigenetic compounds, revealing that histone demethylase inhibitors (HDMi, GSK-J4), with inhibitory effects on multiple lysine demethylases (KDMs)^19^, elicited a distinct repression across all models (Figure 1B). Transcriptional profiling of KDM gene families demonstrated that KDM3 (*KDM3A/B*) and KDM5 (*KDM5A/B*) were the most consistently upregulated lysine demethylases in tumor tissues across intrahepatic CCA patient cohorts (Andersen, TCGA, Job, TIGER^18,20,21^; Figure 1C and S1B; Table S1B), whereas in extrahepatic CCA (Montal cohort^9^), *KDM3B* was the only significantly augmented gene (p=0.04) (Figure 1C and S1C; Table S1B). Overall, these genes did not differ across the main driver mutation subgroups (Figure S1D), suggesting that their deregulation is mutually exclusive from the mutational background.

**Figure 1.**
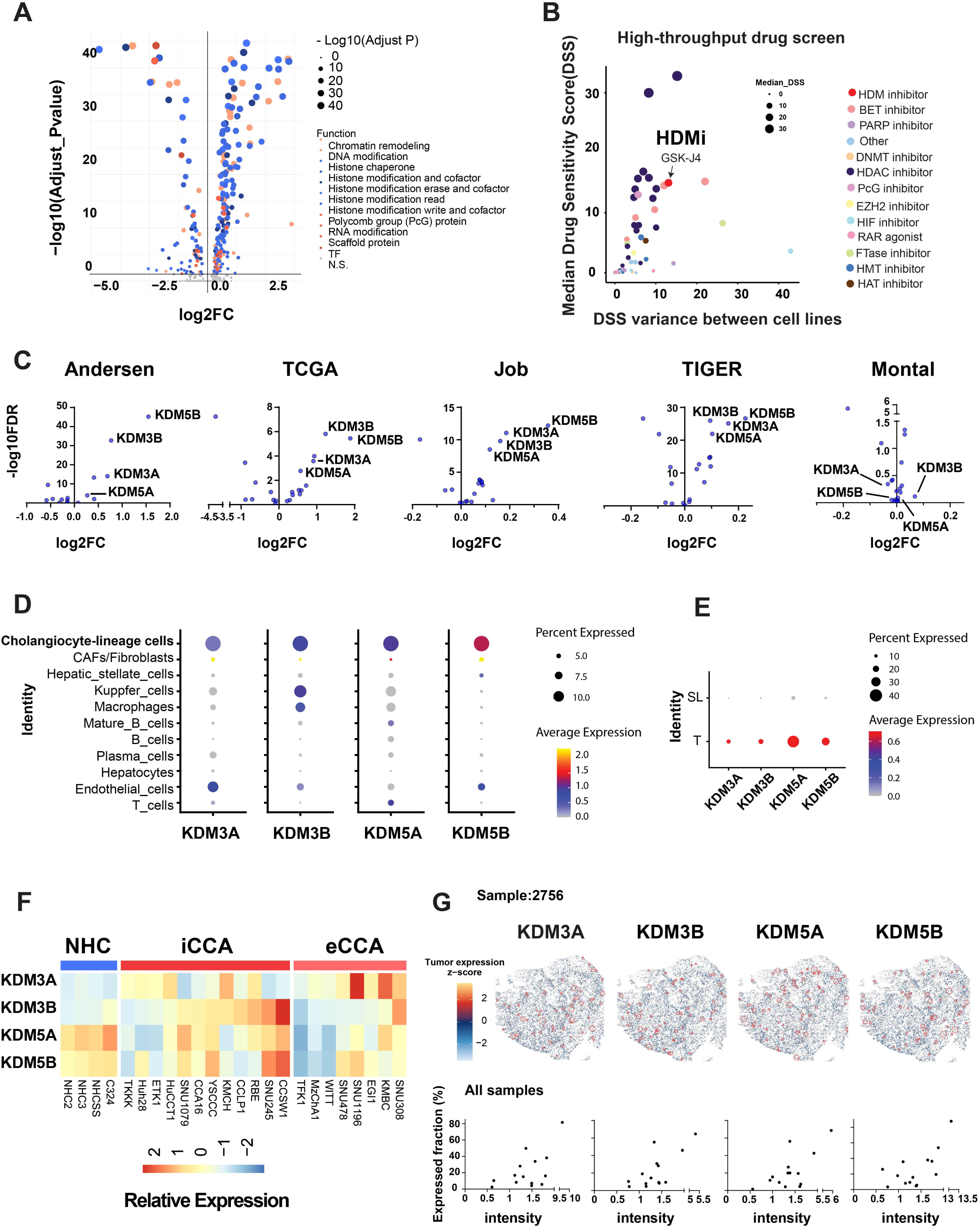
Upregulation of KDM family members in cholangiocarcinoma. (A) Expression of epigenetic factors in patient tumor versus normal tissues in Andersen cohort (n=204 patients). (B) Drug sensitivity scores across eight CCA cell lines from a compound screen. A total of 61 epigenetic compounds were tested. HDMi denotes histone demethylase inhibitor. (C) Expression of all lysine demethylases (KDMs) in tumor tissue compared to surrounding liver across five CCA cohorts (iCCA cohorts: Andersen, TCGA, Job, TIGER, eCCA cohort: Montal). n=415 patients in iCCA and n=182 in eCCA. (D) Expression and fraction of *KDM3A-*, *KDM3B-*, *KDM5A-*, *KDM5B-*expressing cells across cell types in patient scRNA-seq data. (E) Expression of *KDM3A/B* and *KDM5A/B* between tumor (T) and surrounding liver (SL) in cholangiocyte-lineage compartment. (F) Transcriptomic profiling of *KDM3A/B* and *KDM5A/B* in primary normal human cholangiocytes (NHCs), intrahepatic (iCCA) and extrahepatic (eCCA) cholangiocarcinoma cell lines. (G) Spatial feature plots (upper panels) and quantitative summaries (lower panels) depict the expression of *KDM3A*, *KDM3B*, *KDM5A* and *KDM5B* across tumor regions in 14 independent cholangiocarcinoma samples, as assessed by spatial transcriptomics. Red circles indicate merged tumor hotspots, representing contiguous regions enriched for cells with high expression of the indicated gene. See also Figure S1 and Table S1.

To capture the cellular heterogeneity within bulk tumors, we analyzed single cell RNA-seq data from 27 CCA patients, covering in total 44,761 cells obtained from tumor (T) and surrounding liver (SL) tissues (Table S1C)^22–25^. Cell type annotation revealed that *KDM3A/B* and *KDM5A/B* were highly enriched transcripts in the cholangiocyte lineage, encompassing both normal cholangiocytes and tumor cells (Figure 1D and S1E-F). The ductular cholangiocyte cell compartment showed both significantly higher mean expression levels and a higher fraction of *KDM3/5*-positive cells in tumors compared to surrounding livers (Figure 1E). In established CCA cell models, *KDM3A* and *KDM3B* were found to be frequently upregulated relative to primary normal human cholangiocytes (NHCs), whereas this deregulation was not observed for *KDM*5 genes (Figure 1F and Table S1D). Besides, the increased KDM3 expression levels cannot be explained by copy number gains (DepMap data, Figure S1G). Moreover, immunohistochemical validation of KDM expression showed higher KDM3B and KDM5B protein levels in a substantial fraction of CCA tumors (67% for KDM3B and 50% for KDM5B; Human Protein Atlas^12^, Figure S1H), supporting a functional relevance of the augmented KDM regulation.

To examine the distribution of these four genes within patient tumors, we analyzed internal spatial transcriptomics using the Visium HD Spatial Gene Expression platform with 338,427 captured segmented cells from 14 independent intrahepatic CCA specimens^26^ (Table S1E). All four genes displayed regional heterogeneity, with recurrent enrichment across tumor cores, and showed within individual samples coexistence of unique tumor areas presenting heterotopic KDM expression (Figure 1G, upper panel). Expression intensities and the fraction of KDM-positive spatial areas varied substantially between individual tumors (Figure 1G, lower panel). However, the spatial distribution revealed a coordinated regional upregulation and intertumoral variability of the KDM genes rather than showing uniform tumor activation (Figure 1G, upper panel, red circles). As such, the integrated omics and drug screening datasets collectively establish KDMs, particularly KDM3B, as prominently deregulated epigenetic regulators and potential tumor dependencies.

### Inactivation of *KDM3 and KDM5* gene families impairs tumor-like behavior

To investigate the biological role of KDM3/5, we generated knockout (KO) models using CRISPR-Cas9 genome-editing of all four genes in two different CCA models, representing either intra-(HUCCT-1) or extrahepatic (EGI-1) malignancies. Western blotting confirmed a complete loss of each gene in these models prior to assessing several functional phenotypes (Figure 2A, Figure S2A).

**Figure 2.**
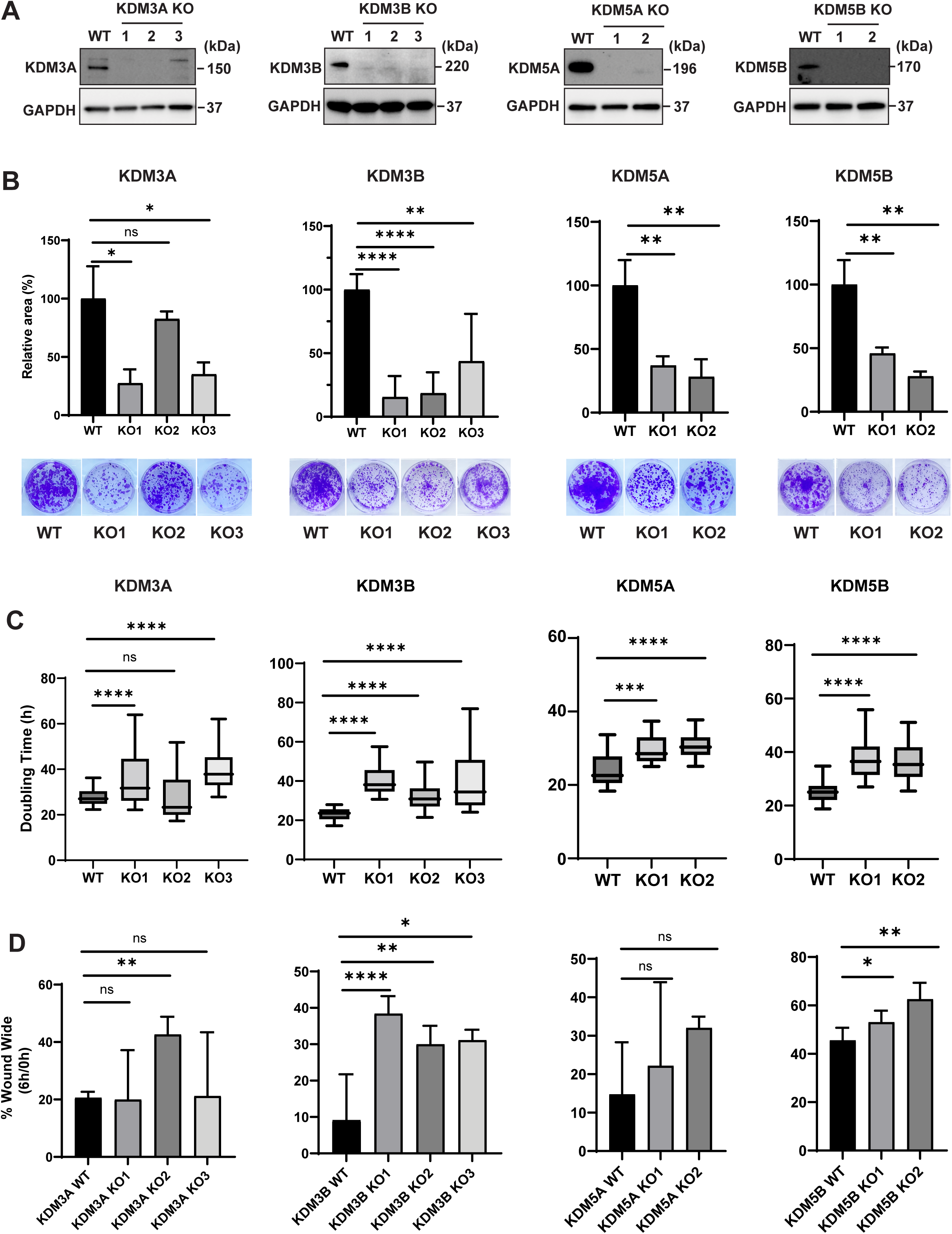
Inactivation of KDM3 and 5 family impairs tumor phenotypes in HUCCT-1 iCCA cell model. (A) The protein of KDM3A, KDM3B, KDM5A and KDM5B were depleted in the knockout (KO) clones by Western blot. GAPDH was used as reference. n=3 independent experiments per condition. (B) Colony formation ability decreased in multiple clones of KDM3 and KDM5 KO by Crystal violet staining. n=3 independent experiments per condition. (C) Doubling time of *KDM3A*, *KDM3B*, *KDM5A* and *KDM5B* KO cells and control (WT). n=2 independent experiments per condition. For each independent experiment, doubling time was calculated from more than 16 consecutive 4-hour measurement intervals during exponential growth. (D) Cell migration was assessed using a wound healing assay. Wound width at 6h was compared with the initial wound width at 0h. n≥3 independent experiments per condition. Mean ± SD are presented with *p* value using unpaired two-tailed Student’s t test (B), (C) and (D); ns, not significant; \**p* < 0.05, \*\**p* < 0.01, \*\*\**p* < 0.001, \*\*\*\**p* < 0.0001. See also Figure S2-3 and Table S2.

Intrahepatic *KDM3B^-/-^*, *KDM5A^-/-^*, and *KDM5B^-/-^*models represented by multiple clones each consistently showed a significant reduction in their ability to form colonies, as well as reduced proliferation with significantly prolonged doubling times across clones, compared to their respective controls (WT) (Figure 2B-C). In contrast, colony formation and proliferation following *KDM3A* loss were more variable between individual clones (Figure 2B-C). Next, evaluating each model’s migratory potential demonstrated an impaired capacity of *KDM3B^-/-^*and *KDM5B^-/-^*cells in closing the gap in the wound healing assay, while there was either no effect (*KDM5A* loss) or mixed results among independent *KDM3A*^-/-^clones on cell migration (Figure 2D).

To evaluate the observed phenotypes in extrahepatic disease, *KDM* genes were comparably deleted in EGI-1 cells, obtaining *KDM3A^-/-^*, *KDM3B^-/-^*, and *KDM5B^-/-^*models. Phenotypically, *KDM3B* loss was consistent with that observed in intrahepatic CCA cells, showing markedly reduced colony formation, proliferation, and migration (Figure S2B-D). In *KDM5B^-/-^*extrahepatic CCA cells, colony formation and the proliferation rate were reduced, whereas the migratory potential remained unaffected. Notably, the effects of *KDM3A* loss differed markedly between model cell lines, with *KDM3A*^-/-^suppressing tumor-like behaviors in HUCCT-1, whereas its loss in EGI-1 cells enhanced colony formation and proliferation, but reduced migration (Figure S2B-D). Together, the assessment of KDM impairment in CCA models revealed that specifically KDM3B exerts consistent and significant impact on tumor cell maintenance by regulating tumor-promoting phenotypes (Table S2), which also corresponds with the assessment of KDM deregulation in patient cohorts.

### *KDM3B* depletion rewires CCA cells

Through phenotypic assessments, we selected KDM3B as the most consistently pro-oncogenic factor (Table S2). To delineate its molecular function, we performed integrated RNA-sequencing, global proteomics, and phosphoproteomics on the HUCCT-1 model using three independent *KDM3B*^-/-^clones (KO1-3) and parallel WT controls. Transcriptomics identified 2,273 genes consistently dysregulated across KO clones (1,162 upregulated; 1,111 downregulated) (Figure 3A and Table S3A-B). As expected, *KDM3B* expression was markedly reduced (padj=1.87×10⁻⁵), whereas other KDMs (such as *KDM3A*, *KDM3C*, *KDM4A-D*, *PHF8*) with overlapping H3K9me1/me2 substrate showed no redundancy (padj>0.05; Figure 3B and Table S3C). These findings were confirmed by qRT-PCR and immunoblotting against the direct KDM3 family (KDM3A and KDM3C) (Figure S3A-B).

**Figure 3.**
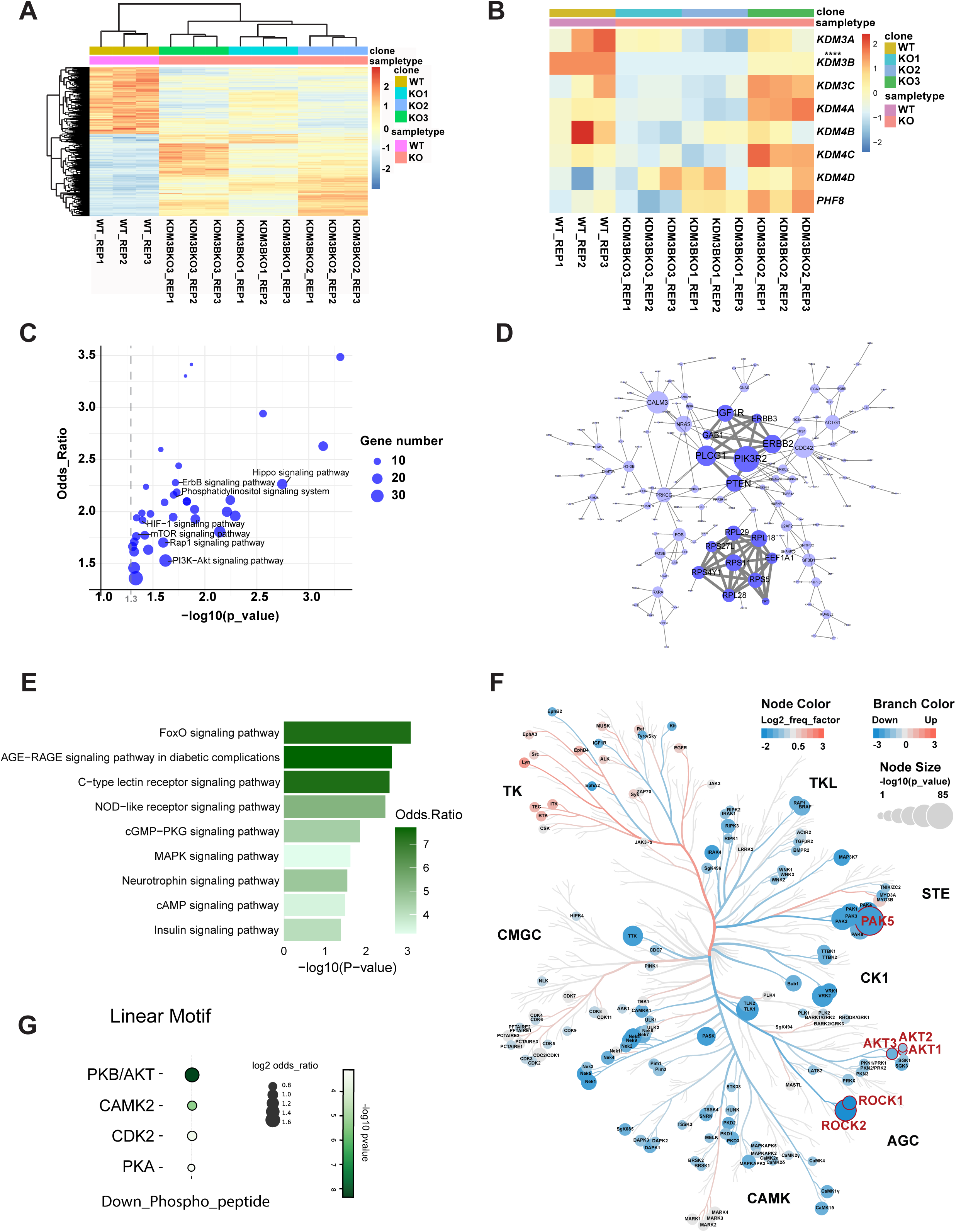
KDM3B depletion suppresses cell growth signaling in CCA cells. (A) Heatmap of significant differential genes (DEGs) between wildtype (WT) and *KDM3B* knockout clones (KO1, KO2 and KO3). (B) Expression of genes with H3K9me1/me2 demethylase activity similar to KDM3B. Normalized RNA-seq read counts were shown across samples. (C) KEGG-defined enriched gene sets (p<0.05) among downregulated genes in *KDM3B*^-/-^cells. Key signaling pathways were highlighted. (D) Protein-protein interaction (PPI) network highlighting core genes among downregulated genes after *KDM3B* loss. (E) KEGG pathway enrichment of downregulated proteins in *KDM3B*^-/-^versus WT cells, identified by whole-proteome LC-MS/MS. (F) Kinome tree showing differential kinase activity between *KDM3B*^-/-^ and WT cells. Significant kinase activity changes (adjusted p<0.05) were identified by phosphoproteomics, with non-significant ones hidden for clarity. (G) Linear motif analysis of down-regulated phospho-targets in *KDM3B^-/-^*versus WT cells by phosphoproteomics. The significant enriched motifs (FDR<0.05) were shown. See also Figure S4 and Table S3.

Given that KDM3B functions as an H3K9me1/me2 demethylase, removing repressive histone marks, its direct transcriptional targets are expected to be downregulated upon *KDM3B* loss. Accordingly, subsequent enrichment analyses were focused on differentially downregulated gene sets, showing that significantly reduced genes are enriched in cell growth-associated signaling networks, including ErbB, Hippo, and PI3K-Akt pathways (Figure 3C and Table S3D). Conversely, upregulated genes are enriched in inflammatory pathways, such as TNF and NFκB signaling (Figure S3C and Table S3E). Protein-protein interaction (PPI) mapping highlighted two major downregulated clusters, including the ErbB-PI3K axis (PIK3R2, PLCG1, ERBB2, IGF1R, PTEN, ERBB3, and GAB1) and a ribosomal protein network (RPS and RPL families) (Figure 3D). Conversely, upregulated PPI networks were enriched for mitochondrial metabolism and TNF-NFκB signaling (Figure S3D).

Next, global proteomics identified 134 upregulated and 111 downregulated proteins consistently across the KO clones, with KDM3B among the most significantly downregulated proteins, confirming effective protein depletion (Figure S3E, Table S3F). Pathway enrichment analysis revealed a significant enrichment in cell growth-related pathways, including FoxO, MAPK, and insulin signaling pathways (Figure 3E, Table S3G), matching the phenotypic assays. Moreover, phosphoproteomics revealed both upregulated (orange) and downregulated (blue) kinase activities, with a prevalent downregulation in phosphorylated sites enriched for STE (MAPK cascade components), AGC (AKT, PKC, and ROCK families), and CAMK (AMPK and calcium-dependent kinases, such as CaMKII) families (Figure 3F and Table S3H) in KO clones, including a pronounced drop in kinase activities affecting cell motility and survival (such as ROCK1, ROCK2 and PAK5). In the KO clones, AKT1-3 exhibited markedly reduced kinase activities, indicating a broad suppression of the PI3K-Akt signaling axis (Figure 3F and Table S3H). Also, linear motif analysis of downregulated phosphoproteins showed significant enrichment of PKB/AKT, CAMK2, CDK2, and PKA motifs, further emphasizing an impairment of the PI3K/Akt signaling and calcium-dependent kinase activity following *KDM3B* loss (Figure 3G). These analyses revealed that KDM3B modulates oncogenic signaling networks, foremost by controlling ErbB and PI3K-Akt signaling pathways, underscoring its central role in sustaining tumor cell maintenance and growth.

### KDM3B controls ErbB signaling

To investigate the regulatory mechanism of KDM3B, we performed chromatin immunoprecipitation sequencing (ChIP-seq) on KDM3B itself, the KDM3B catalytic substrates (H3K9me1 and H3K9me2), and the active histone marks (H3K4me3 and H3K27ac). Peaks representing H3K4me3, H3K27ac, and KDM3B-binding displayed sharp enrichment at promoter regions (Figure 4A). In comparison, H3K9me1 was enriched throughout the gene bodies, whereas H3K9me2 was more broadly distributed upstream of promoters. In *KDM3B^-/-^*compared to WT models, the KDM3B-binding signals were significantly reduced, accompanied by a corresponding increase in H3K9me1 and H3K9me2 signals. Global H3K4me3 and H3K27ac profiles remained largely unchanged (Figure 4A). Differential peak analysis revealed upon *KDM3B*^-/-^, a loss of 26,263 unique KDM3B-binding sites, including at 5,412 promoters. Overall, *KDM3B* loss resulted in 5,796 H3K9me2 gains and 4,015 losses. Similarly, H3K9me1 exhibited 8,667 gains and only 324 peaks lost in *KDM3B*^-/-^versus WT matched controls (Figure 4B). Heatmaps of these differential peaks confirmed a reduced KDM3B occupancy with concomitant increases in H3K9me1/me2 after *KDM3B* loss, whereas H3K4me3 and H3K27ac signals remained unchanged (Figure 4B). Motif analysis of KDM3B-bound regions revealed a strong AP-1 complex transcription factor enrichment, including FosL1, FosL2, and JunB (Figure 4C, Table S4A), suggesting an active recruitment of KDM3B to chromatin via AP-1-mediated interactions.

**Figure 4.**
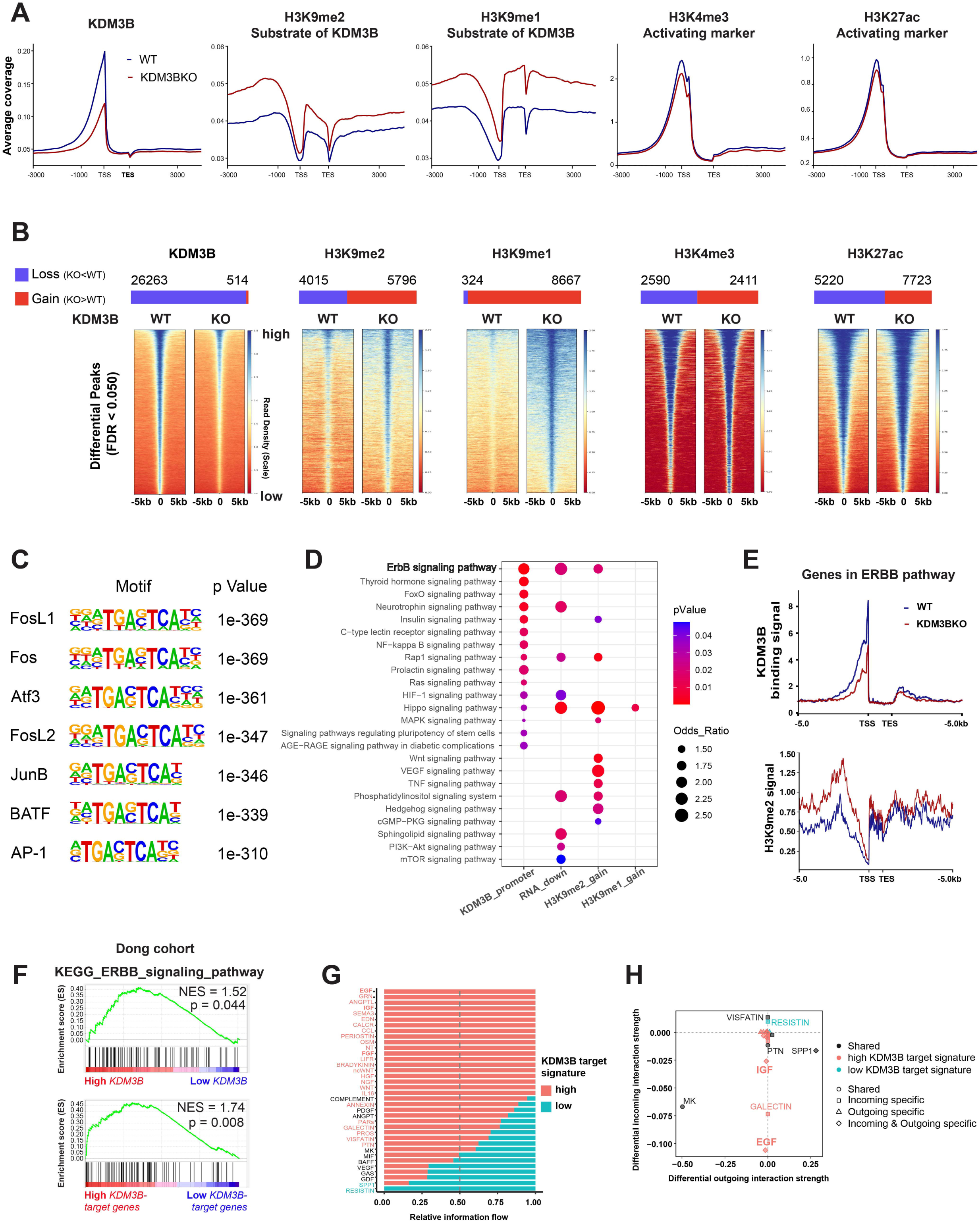
KDM3B is predominantly involved in ErbB signaling regulation. (A) Genome-wide enrichment of KDM3B, H3K9me2, H3K9me1, H3K4me3 and H3K27ac. TSS, transcription start site; TES, transcription end site. (B) Number of differential peaks (upper panel) and heatmaps showing differential peaks occupancy (lower panel) of KDM3B and indicated histone modifications H3K9me1/me2, H3K4me3 and H3K27ac. (C) Top motifs at KDM3B binding sites. *De novo* motif discovery was performed on KDM3B ChIP-seq peaks using *KDM3B*^-/-^ChIP-seq peaks as background. (D) KEGG pathway enrichment of genes with *KDM3B* promoter-binding, decreased RNA expression and increased H3K9me1/2 levels following *KDM3B* loss. (E) Average profiles of KDM3B-binding signal (upper panel) and H3K9me2 signal (lower panel) at genes in ErbB pathway. (F) Gene set enrichment analysis (GSEA) of the ERBB signaling pathway comparing high versus low *KDM3B* expression (upper panel) and KDM3B-target genes (lower panel) in the Dong cohort. (G) Overall information flow of signaling pathways in KDM3B target-gene expression high (top quartile) versus low (lower quartile) groups in tumor cells. Pathways enriched in the KDM3B^high^ group are shown in red, those similarly enriched in both groups are shown in black, and pathways enriched in the KDM3B^low^ group are shown in green. (H) Incoming and outgoing signaling change comparing the high and low KDM3B-target groups in tumor cells. See also Figure S5 and Table S4.

Since KDM3B is a H3K9me1/me2 demethylase, we hypothesized that KDM3B through its catalytic activity directly regulates specific transcriptional programs. To test this, we performed KEGG pathway enrichment analyses, including gene sets showing KDM3B promoter-binding, genes downregulated upon *KDM3B^-/-^*, and genes with increased H3K9me2 or H3K9me1 signals following *KDM3B* loss. This integrated analysis emphasized ErbB signaling as the central pathway controlled by KDM3B, showing a significant reduction in KDM3B-binding and increased H3K9me2 deposition in *KDM3B*^-/-^cells (Figure 4D-E). In total, 36 out of 87 ErbB-related pathway genes showed a significant reduction in KDM3B-binding after knockout, including *AKT3*, *PIK3CA*, *SRC*, *KRAS* (downstream) and *ERBB3* itself, while an additional 34 genes showed a significant increase in H3K9me2 signals (Figure S4A).

By intersection of the three gene sets involving KDM3B promoter-binding, genes showing increased H3K9me2 peaks, and genes with reduced expression upon *KDM3B* loss, we identified 111 genes that significantly displayed promoter occupancy, transcriptional repression, and H3K9me2 accumulation (henceforth the KDM3B target-gene signature; Figure S4B; Table S4B). Confirmatory, we independently showed KDM3B association with ErbB signaling in 255 CCA tumor transcriptomes (Dong cohort^27^), relating that *KDM3B* expression itself and the KDM3B target-gene signature both are positively associated with ErbB signaling (Figure 4F).

Next, applying single-cell transcriptomics on 27 CCA patients^22–25^, we assessed the pathway activity between tumor cells in the top quartile (high) and bottom quartile (low) of the KDM3B target-gene signature. As such, epidermal growth factor (EGF) signaling, the related ligand for pan-ErbB signaling, exhibited the strongest neighborhood ligand-receptor (L-R) interactions, followed by insulin-like growth factor (IGF) signaling (Figure 4G). This interconnectivity suggests that tumor cells with high KDM3B target-gene signature present a strong activation of the ErbB-PI3K-Akt axis, with markedly stronger incoming signals in EGF and IGF activities (Figure 4H). These data establish KDM3B as a central chromatin regulator that through its demethylase role activates the ErbB pathway and a cascade of growth promoting signaling.

### KDM3B directly controls *ERBB3* expression via H3K9me2 regulation

To assess whether KDM3B preferentially regulates specific components of the ErbB signaling, we evaluated individual pathway genes. Thus, we integrated the KDM3B promoter-binding, H3K9me2 peaks, and the DEGs (KO versus WT) showing that *ERBB2*, *ERBB3*, and *CAMK2B* are the top candidate targets of KDM3B (Figure 5A). KDM3B occupied the promoter regions of all three genes. Consistent with KDM3B demethylase activity, *KDM3B* loss resulted in both a significant increase in H3K9me2 enrichment of all three genes and elevated H3K9me1, whereas H3K4me3 and H3K27ac levels remained unchanged (Figure 5B; Table S5A). RNA-seq confirmed that relative *ERBB2*, *ERBB3,* and *CAMK2B* expression levels were significantly decreased in *KDM3B*^-/-^clones compared to controls (Figure 5C). Immunoblotting demonstrated a significant reduction in the ErbB3-encoding HER3 protein level in all clones, with ERBB2 and CAMK2B protein abundance, assessed by mass spectrometry, not exhibiting consistent changes (Figure 5D, Figure S4C).

**Figure 5.**
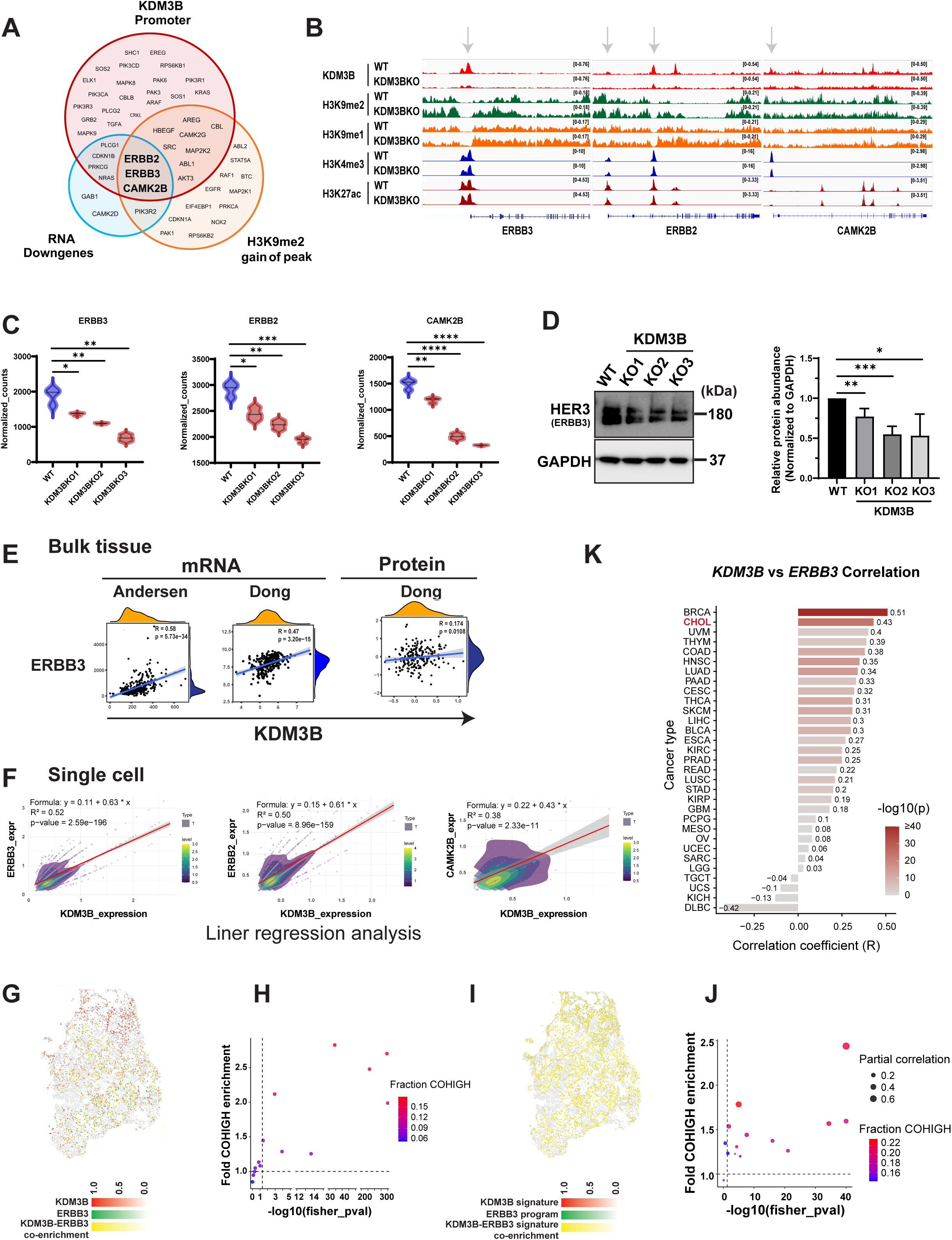
KDM3B regulated the expression of *ERBB3* via H3K9me2 regulation. (A) Overlap of genes in ErbB signaling that are with KDM3B-binding at promoter, H3K9me2 signal increase and RNA down-regulation after *KDM3B* loss. (B) Genome browser view of CHIP-seq signal intensity (KDM3B-, H3K9me1/me2-, H3K4me3- and H3K27ac-occupied regions) at the locus of *ERBB2*, *ERBB3* and *CAMK2B* following *KDM3B*^-/-^ in HUCCT1 cells. (C) RNA level of *ERBB2*, *ERBB3* and *CAMK2B* from RNA-seq following *KDM3B* loss. (D) Protein validation of ERBB3 expressions in *KDM3B^-/-^*models by western blot and quantifications. n=4. Mean ± SD are presented with *p* value using unpaired two-tailed Student’s t test. \**p* < 0.05, \*\**p* < 0.01, \*\*\**p* < 0.001. (E) Association of *KDM3B* and *ERBB3* in RNA (Andersen, Dong cohort) and protein level (Dong cohort) in CCA patients. (F) Association between *KDM3B* expression and *ERBB2*, *ERBB3*, and *CAMK2B* expression in tumor cells revealed by scRNA-seq. (G) Spatial transcriptomic maps show the distribution of KDM3B (red) and ERBB3 (green) expression and their co-enriched spatial spots (yellow) in tumor tissue. (H) Spatial co-enrichment between KDM3B and ERBB3 was quantified across tumor samples (n=14) by fold enrichment of COHIGH spots relative to expectation, with significance assessed by Fisher’s exact test; color denotes the fraction of COHIGH spots per sample. (I) Spatial transcriptomic maps display activity scores of the KDM3B-target gene signature (red), ERBB3 program (green), and spatial spots showing concurrent enrichment of both programs (co-enrichment, yellow). (J) Quantification of spatial co-enrichment between the KDM3B-target gene signature and the ERBB3 program across tumor samples (n=14). (K) Association of RNA expressions of *KDM3B* and *ERBB3* in pan-cancer data from TCGA. See also Figure S5 and Table S5.

PPI hub protein EnrichR-based analysis defined differential peak-associated genes across multiple histone marks (H3K4me3, H3K27ac, H3K9me1, and H3K9me2). Hub genes were identified across all modifications at a significance threshold (p<0.05), with *ERBB2* emerging as the main recurrent hub in the H3K4me3, H3K27ac, and H3K9me2 groups (Figure S4D). *EGFR/ERBB1* and *ERBB3* were also presented as key hubs among the associated hub proteins PPI network, highlighting ERBB signaling members as central nodes among genes with differential histone modifications following *KDM3B*^-/-^(Figure S4E-G). To extend this analysis to CCA patient cohorts, we assessed gene- and protein-level correlations between KDM3B and the central ERBB genes in the Andersen and Dong datasets^5,27^. As such, *ERBB3* expression significantly correlated with *KDM3B* levels at both RNA (R=0.58, p=5.73×10⁻^34^; R=0.47, p=3.20×10⁻¹⁵ in both cohorts, respectively) and at protein levels (R=0.17, p=0.011 Dong cohort) (Figure 5E; Figure S4H). Moreover, linear regression analysis of scRNA data revealed a strong positive association between *KDM3B* and *ERBB3* expression in tumor cells, explaining 52% of the observed variance (R²=0.52, p=2.59×10⁻¹⁹⁶), likewise for *ERBB2* (R²=0.50, p=8.96×10⁻¹⁵⁹) and *CAMK2B* (R²=0.38, p=2.33×10⁻¹¹), supporting their tumor cell-intrinsic relationships (Figure 5F). To determine whether the association between *KDM3B* and *ERBB3* extends pan-cancers, we analyzed TCGA datasets and four additional CCA cohorts^20,21,28,29^, showing a significant correlation in 22 of 31 tumor types (p<0.05), with cholangiocarcinoma (CHOL) at the top (Figure 5K and Table S5B).

Given the strong association between KDM3B and ERBB3 in tumor cells, we evaluated their expression pattern in context of the preserved tumor architecture by spatial transcriptomics. Overlapping signals (yellow) spatially indicate significant co-enrichment of *KDM3B* and *ERBB3* in 8 out of 14 tumor samples (Figure 5G-H). To further evaluate this enrichment at the regulatory level, we compared the KDM3B target-gene signature with ErbB signaling, which revealed significant spatial associations in additionally 12 out of 14 samples (Figure 5I-J). The overlap between *KDM3B* and *ERBB3* at both individual gene and the signature levels support the co-localization of *KDM3B* and *ERBB3* within the same tumor core region. As such, we showed that KDM3B strongly binds to the *ERBB3* promoter and regulates its transcript in a H3K9me2 demethylation-dependent manner. Overall, ErbB signaling genes (*ERBB1-3* and related pathway genes) are generally disturbed by *KDM3B* loss, suggesting an involved cascade effect within this pathway controlled directly by KDM3B bound to *ERBB3*.

### KDM3B-ERBB3 coupling rewires tumor-stromal crosstalk toward a pro-invasive tumor ecosystem

To delineate how KDM3B-ERBB3 co-enriched tumor cells engage with cells in the tumor microenvironment (TME), we performed cell communication analysis in KDM3B-ERBB3 co-high (top quartile) versus co-low (bottom quartile) tumor regions of the 14 intrahepatic CCA specimens (Table S1E). Visualization of signature networks uncovered a qualitative shift in neighborhood signaling. Co-high seed-central neighborhoods were enriched for EMT-associated genes and displayed markedly strengthened tumor-CAF, tumor-endothelial, and tumor-pericyte interactions, with edge weights significantly 4-fold higher than those observed in co-low regions (Figure 6A-B, Table S6A). In contrast, co-low neighborhoods displayed a more heterogeneous signaling enriched for metabolic (glycolysis), angiogenic, and estrogen-response pathways, indicating a negligible stromal/EMT-skewed TME (Figure 6A and Figure S5A, Table S6A). Consistently, tumor cells with top quartile high KDM3B target-gene signature activity displayed increased number of interactions with CAFs compared with tumor cells in the bottom quartile in single-cell RNA analysis (Figure 6C). In the high group, CAFs displayed elevated outgoing interaction strength compared to tumor cells with low KDM3B target-gene signature activity (Figure 6D), supporting that KDM3B^high^ tumor cells enhance stromal engagement. At the cell-type level, directed L-R composition analysis revealed significant differences between tumors in the interactions with CAFs, endothelial cells, pericytes, and myeloid populations, indicating a restructuring of TME communication (Figure 6E). Distinct dominant L-R pairs mediating tumor-neighborhood interactions were observed in co-high and co-low spatial regions within the same tumor tissue, highlighting a rewired tumor-stroma signaling architecture (Figure S5B).

**Figure 6.**
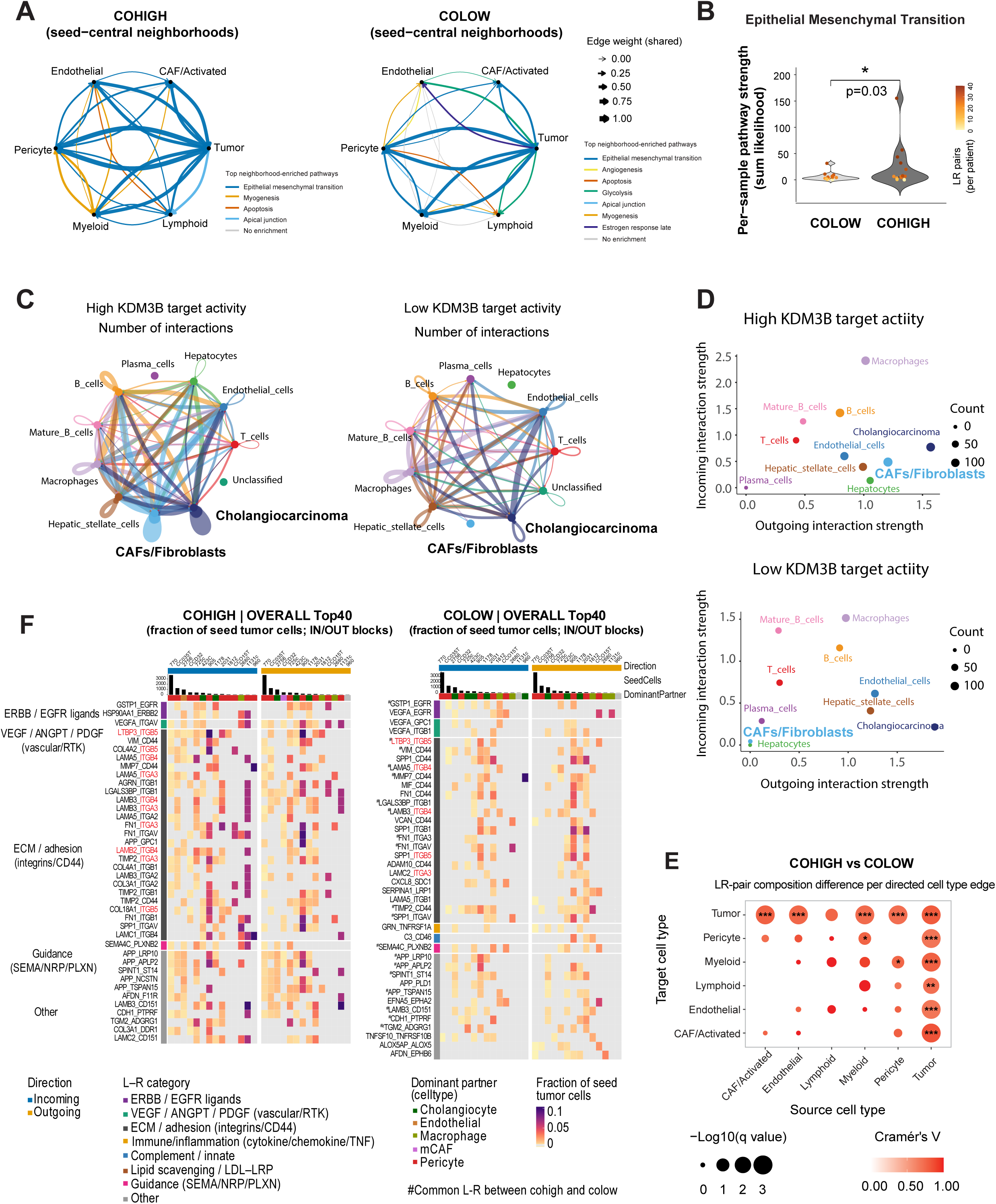
KDM3B-ERBB3 coupling rewires tumor-stromal crosstalk toward a pro-invasive tumor ecosystem. (A) Circle plots depicting intercellular communication networks for the top enriched pathways in KDM3B–ERBB3 co-high and co-low spatial regions. Edge thickness corresponds to interaction strength (aggregated edge weight) between indicated cell types within each pathway. (B) Violin plot showed the strength of co-high and co-low in EMT pathway. Paired-Wilcoxon test, n=13. (C) Number of inferred L–R interactions across cell populations stratified by high versus low KDM3B-target activity (scRNA-seq). (D) Incoming (tumor cells as signal receivers) and outgoing (tumor cells as senders) interaction strengths in high and low KDM3B-target activity groups (scRNA-seq). (E) Comparison of ligand-receptor (L-R) pair composition between co-high and co-low across tumor-neighbor cell-type interactions. Effect size (Cramér’s V) and statistical significance (Monte Carlo–simulated chi-square) are indicated. \**p* < 0.05, \*\**p* < 0.01, \*\*\**p* < 0.001. (F) Heatmap of top 40 L-R interactions between tumor cells and neighboring cell types in KDM3B-ERBB3 co-high or co-low regions within the same tissue section across 14 independent cholangiocarcinoma (CCA) specimens. Ligands or receptors involving KDM3B target genes are highlighted in red. Color intensity indicates the fraction of tumor cells engaged in each interaction. See also Figure S6 and Table S6.

Comparative analysis of the top 40 L-R pairs demonstrated that KDM3B-ERBB3 co-high tumor cells engage more extensively in intercellular communication, with a higher proportion of tumor cells involved in bidirectional signaling across samples, relative to co-low expressing tumor cells (Figure 6F). The fraction of KDM3B-ERBB3 co-high tumor cells exhibited an average of 2.6-fold higher intercellular communication compared to their equivalent co-low counterparts, which preferentially involve ErbB/EGFR ligands, VEGF/ANGPT/PDGF, and extracellular matrix (ECM)/adhesion-associated interactions (Figure 6F, Table S6B). Moreover, integrin-associated (ITGA/B family members) L-R pairs were markedly enriched in the co-high signaling landscape, comprising 25 of the top 40 overall tumor cell incoming signals. Specifically, ECM-adhesion (such as LTBP3-ITGB5 and LAMC1-ITGB4) engaged up to ∼44% and ∼40% of cells in co-high tumors, respectively, whereas the strongest structural interaction in co-low tumors (SPP1-CD44) reached only 24% of the cell fraction (Table S6B). This integrin-mediated adhesion of KDM3B-ERBB3 co-high tumor cells is consistent with the impaired migratory capacity observed upon KDM3B depletion *in vitro* (Figure 2D). Compared to co-low regions, co-high neighborhoods exhibited a higher fraction of pericytes, with pericytes acting at a substantially increased frequency as the dominant interacting partner in co-high signaling networks (Figure 6F and Figure S5C). This is in accordance with the role of pericytes in promoting tumor cell motility and metastatic dissemination^30,31^. Moreover, the top 40 L-R interactions in co-high tumor regions demonstrate a set of uniquely recruited endothelial cells and cancer-associated fibroblasts (CAFs) as dominant tumor cell partners, which were absent in co-low regions, suggesting a rewired and more stromal tumor ecosystem, that engages KDM3B-ERBB3 regulation (Figure 6F). Among these ligand-receptors, *LTBP3*, *LAMB2*, *ITGA3*, *ITGB4*, *ITGB5* were identified within KDM3B-target signature, exhibiting promoter-bound KDM3B, increased H3K9me2 deposition, and reduced RNA expression following KDM3B deletion (Figure 6F, Figure S5D and Table S4B). Specifically, focusing on ERBB3-centered signaling, we observed a relative enrichment of ERBB3-associated L-R pairs in co-high tumors, whereas these interactions are rare in co-low cells (Figure S5E). Collectively, KDM3B-ERBB3 coupling rewires tumor-stromal interactions to promote a pro-invasive tumor ecosystem.

### Targeting the ErbB-Akt axis downstream KDM3B regulation offers therapeutic potential in CCA

To evaluate whether KDM3B dysregulation of the ErbB-Akt signaling presents an option to inhibit CCA tumor growth, we assessed pathway-targets in xenograft models. As KDM3B-specific inhibitors are unavailable and KDMi generally presents unwanted toxicities, we chose first to directly target the ErbB receptors using trastuzumab/T, an approved monoclonal antibody that disrupts the HER2-HER3 (*ERBB* coding proteins) interaction and attenuates ErbB signaling^32^. CCA xenograft tumors were established by injecting HUCCT-1 cells subcutaneously into the flanks of immunocompromised mice (n=55, in total), tumor volumes were measured weekly, and animals were randomized prior to treatment when tumors reached palpable size at approximately 40 mm³ each (trastuzumab study) and larger established tumor volumes at 200 mm^3^ for the later pan-AKTi study (Figure 7A). Trastuzumab treatment (0.45mg/dose i.p.) was administered twice weekly over a 4-week period and observed to significantly reduce tumor volumes compared with control/vehicle (P_effect_<0.02, Figure 7B). At the end of study, tumors were extracted and transcriptomes profiled, establishing a trastuzumab response-signature comprising 110 differentially expressed genes (p<0.001) (Figure 7C). Hierarchical clustering using this gene set stratified CCA patients in both the Andersen and Dong cohorts into two unique groups, with the patient subset enriched in the signature genes representing improved overall survival (OS) (Figure 7D and Figure S6A). These data were validated at the matched protein level in the Dong cohort, where clustering by the corresponding signature separated patients with significant differential outcomes (Figure 7E). Enrichment analysis on the tumor proteomes showed that when comparing clusters separated by the signature genes, patients in cluster1 with shorter survival time, their tumors highly express ErbB signaling and downstream effectors, including insulin, VEGF, and MAPK signaling (Figure S6B).

**Figure 7.**
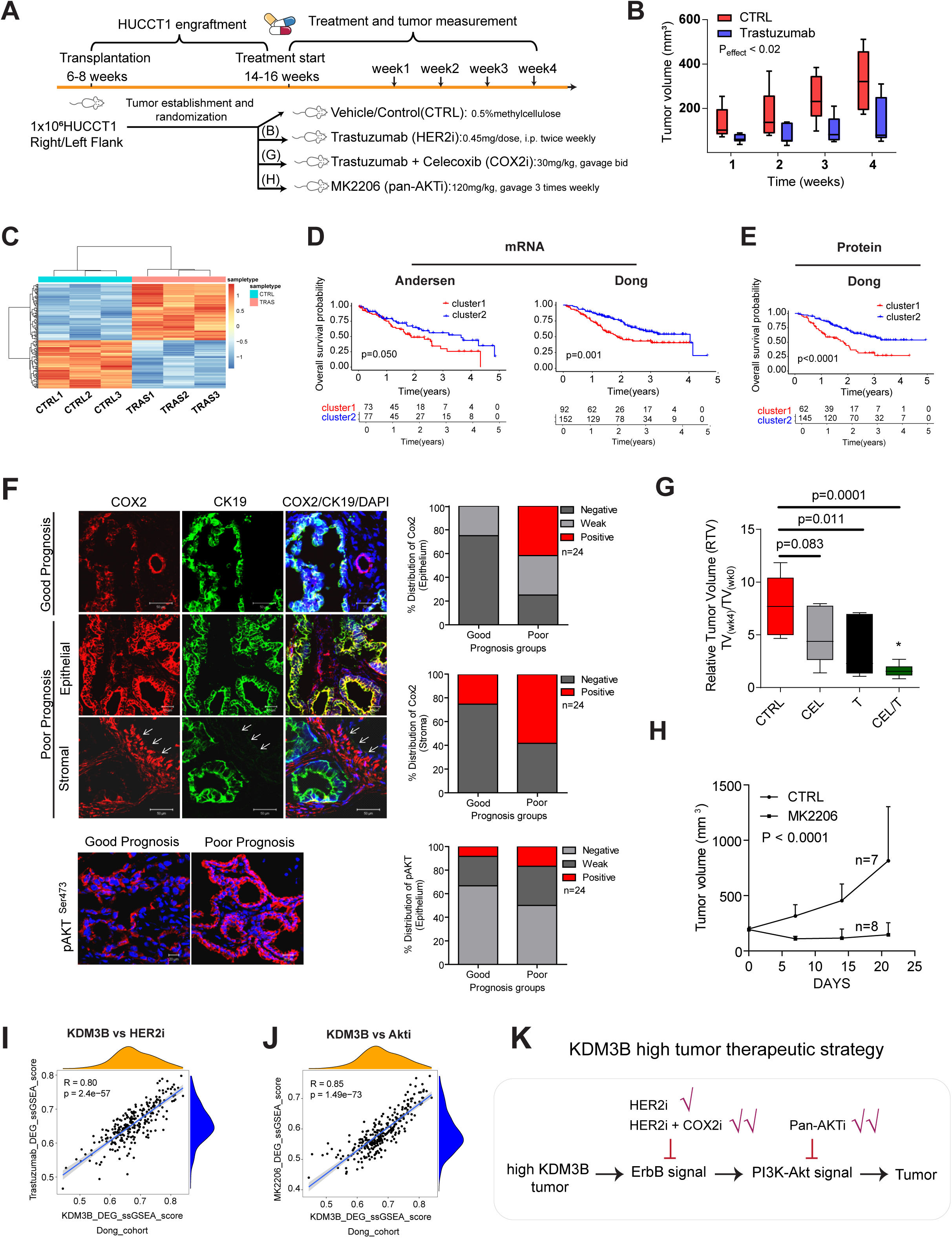
Targeting the KDM3B-ErbB-Akt axis exhibited anti-tumor effect *in vivo*. (A) Illustration of experimental design in CCA xenograft model with multiple treatment in ErbB-PI3K/Akt axis. (B) Quantification of *in vivo* tumors (n≥6 per group) progression via tumor volume over time after the start of the treatment with trastuzumab (HER2 inhibitor). Statistical significance is determined by 2-way ANOVA. (C) Identification of a 165-gene response signature from transcriptomic analysis of DEGs (p<0.001) in trastuzumab-treated versus control tumor tissues. (D) Kaplan-Meier survival curves of CCA patients stratified by trastuzumab response signature at transcriptional level in Andersen and Dong CCA cohorts. (E) Kaplan-Meier survival curves of CCA patients stratified by trastuzumab response signature at protein level in Dong cohort. Log-rank (Mantel–Cox) test for (C) and (D). (F) Immunofluorescence staining of COX2 (red) and tumor marker CK19 (green) in stromal and epithelial compartments of patients with good versus poor prognosis^18^. (G) Relative tumor volume (RTV) in control (CTRL), trastuzumab-treated (T), celecoxib-treated (CEL), and combination-treated groups (CEL/T). Tumor volumes at week 4 were normalized to baseline (week 0, treatment initiation). n≥6 tumors per group. (H) Xenograft tumor volume following treatment with MK2206 (Pan-AKT inhibitor) or vehicle (n ≥ 6 tumors per group). (I) Correlation of ssGSEA scores between KDM3B knockout differential expressed genes (DEGs) and trastuzumab-responsive DEGs in the Dong cohort (n=255). (J) Correlation of ssGSEA scores between KDM3B knockout DEGs and MK2206-responsive DEGs in the Dong cohort. (K) Summary of targetable therapeutic strategy in KDM3B^high^ tumor. See also Figure S7 and Table S7.

Applying univariate Cox regression analysis on the trastuzumab response-signature, we defined a gene subset significantly associated with OS. Validating these genes against 662 tumors (4 individual CCA cohorts)^18,20,27,29^, we unveiled cyclooxygenase-2 (COX2), a superior independent predictor of OS (Figure S6C-D). Immunofluorescence staining of COX2 in tumors from patients stratified based on survival (prior transcriptomic prediction^18^) combined with the epithelial marker cytokeratin 19 (CK19), COX2 was markedly elevated in both epithelial and stromal compartments and significantly enriched in patients with poor survival outcomes (also high in ErbB signaling^18^) (Figure 7F), which supports the role of activated ErbB signaling in relation to COX2, promoting tumor growth. Thus, we hypothesized that blocking COX2 signaling using the nonsteroidal anti-inflammatory drug celecoxib/CEL either alone or in combination with trastuzumab would present an added beneficial effect by reducing the xenograft tumor volumes. Alone, celecoxib (30mg/kg, gavage) was insufficient in reducing the relative tumor volume (RTV), but in combination with trastuzumab, the effect of blocking the ErbB receptor signaling was markedly improved, significantly reducing the xenograft tumor burden over 4 weeks (Figure 7G).

Next, we showed that AKT activation (serine 473 phosphorylation) is significantly higher in the same patient group with poor prognosis (Figure 7F). Thus, we tested the efficacy of pan-AKTi (MK2206) in blocking xenograft tumor growth, showing significantly reduced tumor volumes and tumor weights relative to vehicle-treated mice (Figure 7H, S6E-F). This demonstrates that targeting several steps in the KDM3B-ERBB3 signaling axis presents options in CCA, emphasizing the KDM3B regulation and downstream biology. As such, we examined the relationship between KDM3B and ErbB-AKT signaling by correlating the ssGSEA scores between KDM3B-, trastuzumab-, and MK2206-responsive genes across CCA patient cohorts (Table S7A). This analysis revealed a strong correlation between genes controlled by KDM3B and trastuzumab/pan-AKTi-responsive genes (Figure 7I-J, S6G-H and Table S7B), supporting a significant association between KDM3B histone lysine demethylase activity and downstream ErbB-Akt signaling. Together, these *in vivo* studies demonstrate the therapeutic strategy for *KDM3B* high tumor via ErbB blockade, AKT inhibition, or a combined ERBB-COX2 strategy (Figure 7K).

## Discussion

Epigenetic deregulation is a defining genomic feature of cholangiocarcinoma (CCA)^33^. In this study, we elucidated a previously unrecognized dependency on histone lysine demethylase (KDM) delineating a mechanistic axis through which KDM3B prevalently activates ERBB3-ErbB signaling to reprogram tumor cell signaling and the TME to sustain malignant growth.

Multiple CRISPR-based perturbations in CCA models established *KDM3B* as the dominant gene on cell fitness. Integrated analyses converged on ErbB controlling PI3K-AKT as the principal axis attenuated upon *KDM3B* loss, with direct KDM3B occupancy at the *ERBB3* promoter, gain of repressive H3K9me2, and concordant downregulation of ERBB3 gene and protein levels as well as attenuated ErbB signaling. Patient transcriptomes demonstrated elevated KDM3B activity with associated augmented ErbB signaling and significant ERBB3 tumor cell expression. Spatial transcriptomic and single-cell datasets emphasized that KDM3B cooperates with ERBB3 to reshape intercellular communication networks, reinforcing EMT-associated signaling and strengthening neighborhood cell interactions to establish a supportive TME. Inhibiting the KDM3B-ErbB-Akt signaling effectively reduces xenograft tumor growth. Thus, these findings position KDM3B as an epigenetic gatekeeper of ERBB3-dependent intercellular signaling and intercellular crosstalk, creating a vulnerability in CCA.

Non-mutational epigenetic reprogramming is increasingly recognized as a hallmark of cancer, representing a distinct mechanism that establishes oncogenic chromatin states and promotes tumor development^3,34^. In this setting, we identified KDM3B as a non-mutational epigenetic regulator essential for tumor maintenance in CCA. Multi-omics analyses consistently converged on a central role for KDM3B in sustaining ERBB signaling, a key oncogenic pathway driving CCA development and progression^1,2,18^. *KDM3B* loss attenuated kinase activities associated with the PI3K-AKT, AMPK, and MAPK pathways, indicating broad suppression of ERBB downstream signaling. Mechanistically, ChIP-seq demonstrated direct binding of KDM3B at the *ERBB3* promoter, where KDM3B-mediated H3K9me2 demethylation promotes *ERBB3* transcription. Elevated ERBB3/HER3 may heterodimerize with ERBB2/HER2, thereby activating ErbB signaling and its downstream PI3K-AKT cascade to support tumor growth^18,36–38^. Consistent with this mechanism, CCA patient spatial transcriptomics revealed enhanced EGF- and IGF-mediated intercellular communication in KDM3B^high^ tumor cells, including co-enrichment of *KDM3B* and *ERBB3* within tumor regions. Moreover, KDM3B-regulated transcripts strongly associated with ErbB2 and AKT inhibitor-response genes in patient cohorts, emphasizing the KDM3B-ErbB-Akt regulatory axis. Also, these findings are supported by previous observations that *Kdm3b* deficiency impairs somatic growth in mice through reduced IGF1 signaling^39^.

Spatial transcriptomics further revealed that the KDM3B-ERBB3 axis reshapes ligand-receptor communication between tumor cells and the surrounding stromal TME. Tumors with KDM3B^high^ activity exhibited enhanced EMT-associated crosstalk with CAFs, endothelial cells, and pericytes, together with strengthened ECM-integrin interactions. This enhanced tumor-stroma communication, particularly through ECM-integrin signaling, is known to promote EMT and establish a more pro-invasive tumor niche^40^. Several ECM-related ligand-receptor components were identified as direct transcriptional targets of KDM3B, suggesting that the remodeling of the tumor microenvironment is driven not only by enhanced ERBB signaling^41^ but also by direct rewiring of the ECM-integrin signaling network. Increased ECM-integrin engagement may activate the FAK/SRC signaling axis, which has been shown to crosstalk with ErbB signaling^42,43^. Thus, KDM3B-driven ECM remodeling, ERBB activation, and EMT collectively establish a positive feedback loop that promotes a pro-invasive tumor microenvironment and supports tumor growth.

The functional coupling between KDM3B and ERBB3 highlights a potential therapeutic vulnerability downstream of KDM3B. Pharmacological inhibition of HER2 (trastuzumab), pan-AKT, and combined HER2/COX2 targeting, each suppressed tumor growth. Although epigenetic therapies have shown limited efficacy as monotherapies in solid tumors owing to insufficient selectivity and dose-limiting toxicities, accumulating evidence suggests that combination strategies may offer greater clinical benefit^44–47^. Our findings provide a rationale for which KDM3B inhibition may potentially enhance ErbB-PI3K-AKT pathway inhibition. Currently, HER2-targeted therapies in cholangiocarcinoma, such as zanidatamab, are approved against IHC3+ HER2-positive tumors. Since KDM3B transcriptionally activates ERBB3 and sustains ERBB signaling independently of HER2 amplification, our results raise the possibility that tumors with KDM3B^high^ activity may also benefit from such drugs, likely expanding the target population.

Our study is limited by the absence of a selective KDM3B inhibitor. Some inhibitors (P3FI-90, GSK-J4) have binding and inhibitory effects on KDM3B, but their specificities are limited by also targeting other lysine demethylases^48,49^. Pharmacologic confirmation will require future structure-guided ligands or PROTACs exploiting non-catalytic KDM3B surfaces for selectivity. Second, although xenograft models can capture direct on-target pathway effects, patient-derived organoids or genetically engineered CCA models could refine further treatment strategies since not all tumors may rely on the KDM3B→ERBB3/ErbB axis^34^.

## RESOURCE AVAILABILITY

### Lead contact

Further information and requests for resources and reagents should be directed to and will be fulfilled by the lead contact, Jesper B. Andersen.

### Material, Data and Code availability

The RNA-seq and CHIP-seq raw and processed data have been deposited in the Gene Expression Omnibus (GEO) under accession numbers: GSE313018 and GSE313100, respectively. The microarray raw and processed data have been deposited in the GEO under accession numbers: GSE342162. Global proteomic and phosphoproteomic datasets have been deposited in the ProteomeXchange Consortium via PRIDE under accession numbers PXD071760 and PXD071761. This paper also analyzes existing, publicly available datasets (accession number and link are found in the key resources table). This paper does not report original code. Any additional information required to reanalyze the data reported in this paper or materials are available from the lead contact upon request. Further information can be found online.

## ACKNOWLEDGEMENT

The authors are grateful to Prof. Snorri S. Thorgeirsson, Drs. Valentina Factor and Elizabeth A. Conner from the National Cancer Institute, NIH in Bethesda, USA for help with the *in vivo* modeling. Visium HD work was supported by CellX (The Danish Single Cell Examination Platform), funded by the Danish Research Agency through the Danish national research infrastructure program (5229-0009B). Mass spectrometry-based proteomics analyses were performed at the Proteomics Research Infrastructure (PRI), University of Copenhagen (UCPH), supported by the Novo Nordisk Foundation (NNF) (grant agreement number NNF19SA0059305). We thank for technical help Irina Korshunova, Bente Emma Møller, and Eman Ahmad Mouhammad in BRIC Single Cell Genomics Core Facility. CCA tumor tissues were originally provided by Prof. Lewis R. Roberts supported in parts by Award Number P50 CA210964 (Mayo Clinic Hepatobiliary SPORE) from the National Cancer Institute (NCI), USA. The content is solely the responsibility of the authors and does not necessarily represent the official views of the NIH, USA. The authors thank Drs. Leonor Rib, Yasuko Antoku, André Eduardo Carneiro Dias, Elin Josefina Pietras from the BRIC High Content screening and Microscopy core facilities utilized during this study. The results are in part supported by data generated by the TCGA Research Network: https://www.cancer.gov/tcga.

## Author contributions

Conception and design: J.B.A., C.J.O., Y.L.; methodology: Y.L., J.B.A., S.L., A.M., A.T., S.Y., M.W.; meta-analysis: Y.L., O.N., E.Z., C.J.O, A.M., C.N., A.R.C.; interpretation of data: Y.L., J.B.A., O.N., C.J.O., J.L.B., C.N.; patient omics and clinical data collection: J.B.A., C.J.O., E.Z., C.N., Y.L., S.Y.; visualization: Y.L., O.N., E.Z., J.L.B., C.N., A.R.C., J.B.A; writing-original draft: Y.L., J.B.A., O.N.; All authors read, edited, and approved the manuscript.

## STAR METHODS

- KEY RESOURCES TABLE
- RESOURCE AVAILABILITY

`o` Lead contact
`o` Materials availability
`o` Data and code availability
- EXPERIMENTAL MODEL AND STUDY PARTICIPANT DETAILS

`o` Mouse models
`o` Cell lines
- METHOD DETAILS

`o` Generation of knockout cell lines
`o` Epi-drugs sensitivity analysis
`o` Expression analysis of CCA cell lines
`o` Chromatin immunoprecipitation sequencing (ChIP-seq)
`o` Immunoblotting
`o` Reverse-transcriptase quantitative PCR (RT-qPCR)
`o` Colony formation assay
`o` Cell proliferation
`o` Wound healing assay
`o` Transcriptomics
`o` Immunofluorescence
`o` Global proteomics and Phosphoproteomics
`o` Patient characterization and omic datasets
`o` RNA-seq analysis
`o` CHIP-seq analysis
`o` Proteomic and Phosphoproteomic analysis
`o` Single-cell RNA-seq (scRNA-seq) analysis
`o` Gene dependency analysis
`o` Copy number alteration analysis
`o` Protein-protein interaction (PPI) network analysis
`o` Gene Set Enrichment Analysis (GSEA)
`o` Gene and gene set association in patient dataset
`o` Visium HD spatial transcriptomic data processing
`o` *KDM3B-ERBB3* co-distribution analysis
`o` Spatially ligand-receptor and cell-cell communication analysis
- QUANTIFICATION AND STATISTICAL ANALYSES

## Supplemental table titles

**Table S1A.** Dysregulated epigenetic factors in CCA tumor tissue compared with surrounding liver and bile ducts. Related to Figure 1.

**Table S1B.** Clinical characteristics of CCA cohorts used in this study. Related to Figure 1, 4-6.

**Table S1C.** scRNA data sets and information in this study. Related to Figure 1, 4-5.

**Table S1D.** RNA expression of *KDM3A/B* and *KDM5A/B* in normal and cholangiocarcinoma cell lines. Related to Figure 1.

**Table S1E.** Clinical information of VISIUM HD spatial transcriptomics samples. Related to Figure 1, 5 and 6.

**Table S2.** Evaluation of phenotypes following KDM3A/B and KDM5A/B knockout. Related to Figure 2 and Figure S2.

**Table S3A.** Normalized counts in KDM3B knockout versus wild-type. Related to Figure 3.

**Table S3B.** Genes consistently upregulated or downregulated across three KDM3B knockout clones versus wild-type. Related to Figure 3.

**Table S3C.** Expression of potential compensatory genes following KDM3B depletion. Related to Figure 3.

**Table S3D.** KEGG enrichment of downregulated genes after KDM3B knockout. Related to Figure 3.

**Table S3E.** KEGG enrichment of upregulated genes after KDM3B knockout. Related to Figure 3.

**Table S3F.** Differentially expressed proteins in global proteomics of KDM3B knockout clones (KO1, KO2, KO3) compared with wild-type. Related to Figure 3.

**Table S3G.** KEGG enrichment of globally downregulated proteins. Related to Figure 3.

**Table S3H.** List of kinases with significantly altered activity upon KDM3B knockout. Related to Figure 3.

**Table S4A.** Homer known motifs related to KDM3B binding sites. Related to Figure 4.

**Table S4B.** KDM3B related gene sets. Related to Figure 4.

**Table S5A.** Differential signals of KDM3B, H3K9me1, and H3K9me2 in the gene region of ERBB2, ERBB3, and CAMK2B. Related to Figure 5.

**Table S5B.** Correlation between KDM3B and ERBB3 in pan-cancer transcriptomic data from patient tissues in TCGA. Related to Figure 5.

**Table S5C.** Gene list of ERBB3 program. Related to Figure 5.

**Table S6A.** Interaction strength in the top enriched pathways. Related to Figure 6.

**Table S6B.** Cell fractions engaged in ligand-receptor-mediated intercellular communication. Related to Figure 6.

**Table S7A.** Gene sets for signature association. Related to Figure 7.

**Table S7B.** ssGSEA scores of different gene sets in transcriptomic data from the Dong and Andersen CCA cohorts. Related to Figure 7.

**Table S8A.** List of sgRNA sequences used for cloning, including BsmBI recognition site. Related to STAR Methods.

**Table S8B.** Primers used in the study. Related to STAR Methods.

