## supplementary figures and material and method for "KDM3B enforces a non-mutational ErbB signaling state that sustains cholangiocarcinoma"

#### This PDF file includes:

Supplementary Figures S1 to S6

Figure legends of Figures S1 to S6

Materials and Methods

Key resources table

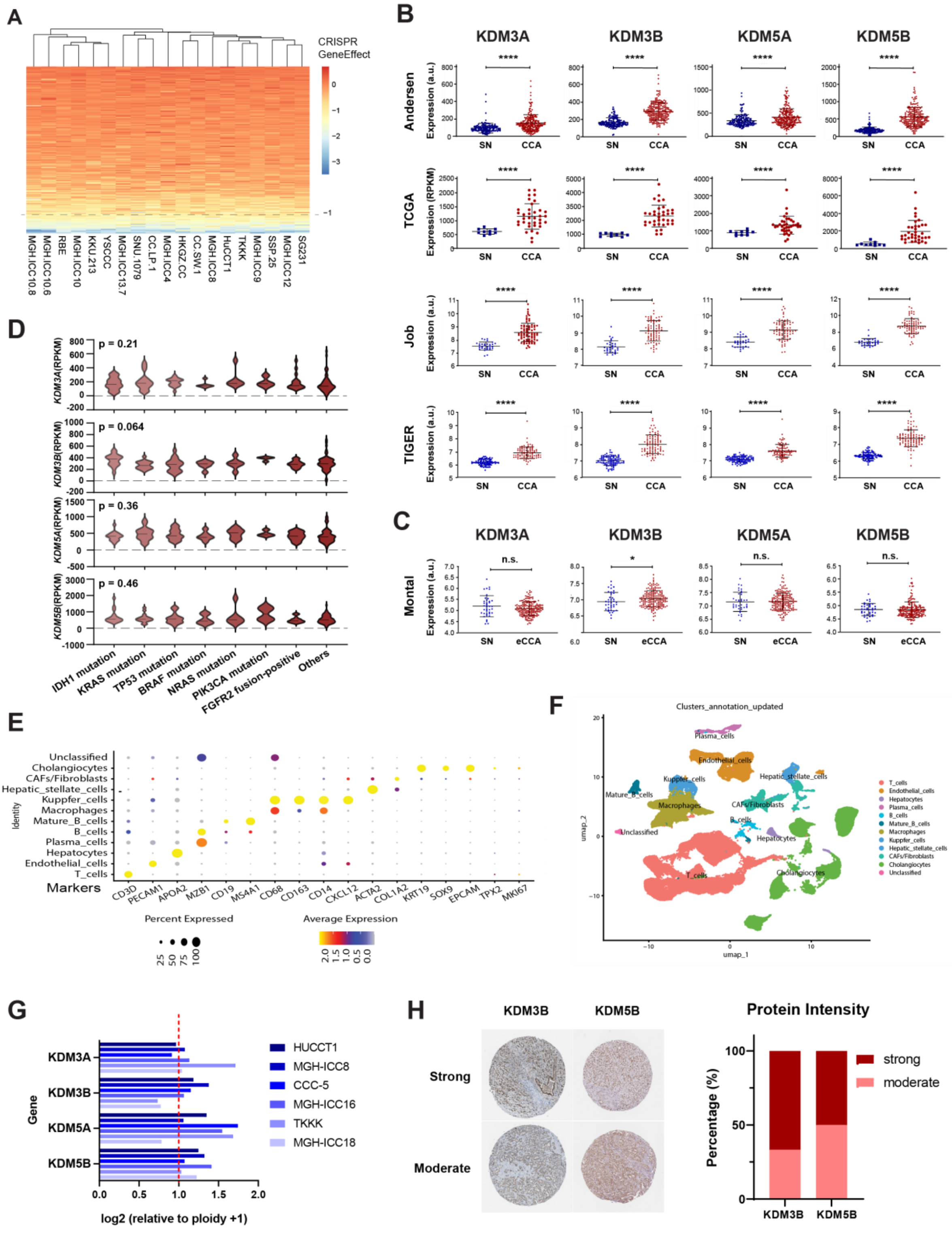
A
CRISPR GeneEffect
MGH.ICC10.8
MGH.ICC10.6
RBE
MGH.ICC10
KKU.213
YSCCC
SNU.1079
MGH.ICC13.7
CC.LP.1
MGH.ICC4
HKGZ.CC
CC.SW.1
MGH.ICC8
HuCCT1
TKKK
MGH.ICC9
SSP.25
MGH.ICC12
SG231
B
KDM3A
KDM3B
KDM5A
KDM5B
Andersen
TCGA
Job
TIGER
Expression (a.u.)
Expression (RPKM)
SN
CCA
C
Montal
n.s.
SN
eCCA
D
p = 0.21
p = 0.064
p = 0.36
p = 0.46
KDM3A(RPKM)
KDM3B(RPKM)
KDM5A(RPKM)
KDM5B(RPKM)
IDH1 mutation
KRAS mutation
TP53 mutation
BRAF mutation
NRAS mutation
PIK3CA mutation
FGFR2 fusion-positive
Others
E
Unclassified
Cholangiocytes
CAFs/Fibroblasts
Hepatic_stellate_cells
Kupffer_cells
Macrophages
Mature_B_cells
B_cells
Plasma_cells
Hepatocytes
Endothelial_cells
T_cells
Identity
Markers
CD3D
PECAM1
APOA2
MZB1
CD19
MS4A1
CD68
CD163
CD14
CXCL12
ACTA2
COL1A2
KRT19
SOX9
EPCAM
TPX2
MKI67
Percent Expressed
Average Expression
F
Clusters_annotation_updated
umap_1
umap_2
G
KDM3A
KDM3B
KDM5A
KDM5B
Gene
HUCCT1
MGH-ICC8
CCC-5
MGH-ICC16
TKKK
MGH-ICC18
log2 (relative to ploidy +1)
H
KDM3B
KDM5B
Strong
Moderate
Protein Intensity
strong
moderate
Percentage (%)

**Figure S1. Expression of *KDM3A/B* and *KDM5A/B* in human CCA tumor samples, related to Figure 1.** (A) DepMap CRISPR gene-effect scores in CCA cell lines for significantly dysregulated epigenetic regulators identified in Fig. 1A. Scores = 0, no effect on cell viability; Scores < -1, significant dependency required for cancer cell survival. (B) Expression of *KDM3A/B* and *KDM5A/B* in tumors (CCA) versus matched tumor-adjacent surrounding normal (SN) tissue in iCCA cohorts of Andersen, TCGA, Job and TIGER. (C) Expression of *KDM3A/B* and *KDM5A/B* in CCA versus matched SN in eCCA cohort Montal. Statistical significance is determined by Mann-Whitney test in (B) and (C). ns, not significant; $*p < 0.05$, $***p < 0.001$, $****p < 0.0001$. (D) Expression of *KDM3A/B* and *KDM5A/B* in tumor tissues from the Andersen cohort stratified by mutation status. Mutation presents in at least four patients were included in the analysis. Statistical significance was determined by Kruskal-Wallis's test. (E) Feature plots of selected marker genes used for cell-type annotation in single-cell RNA-seq (scRNA) data. (F) UMAP of cell types from scRNA data. (G) Copy numbers of *KDM3A/B* and *KDM5A/B* in CCA cell lines from DepMap. (H) Protein intensity of KDM3B and KDM5B in cholangiocarcinoma tumor tissues via immunohistochemistry (IHC) from Human Protein Atals database[1]. n = 12 for KDM3B and n = 8 for KDM5B.

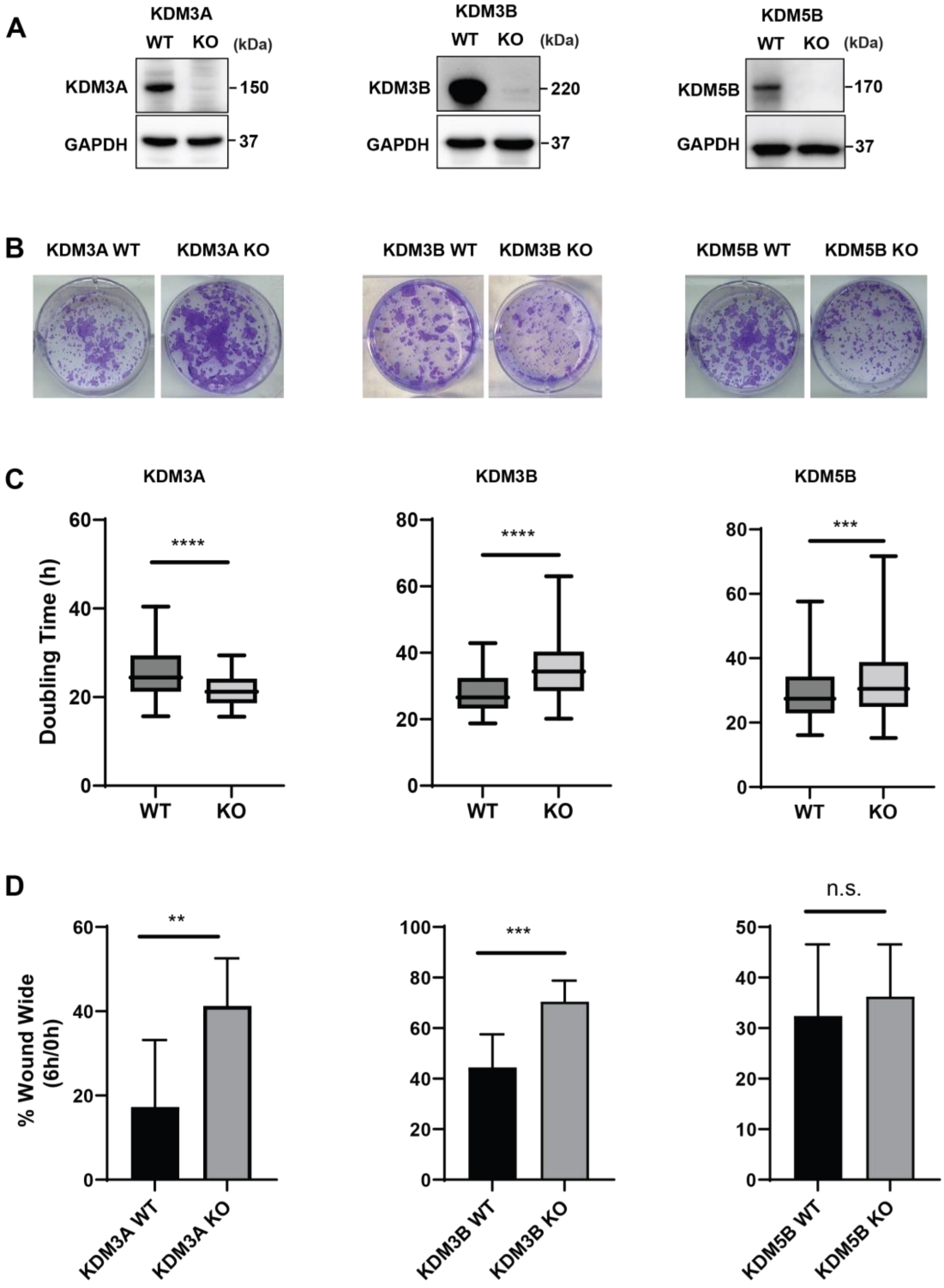

**Figure S2. Characterization of *KDM3A*, *KDM3B* and *KDM5B* knockout in EGI-1 (eCCA) cell models, related to Figure 2.** (A) Protein depletion of KDM3A, KDM3B and KDM5B in the respective knockout clone, confirmed by western blot. GAPDH was used as reference. n=2 independent experiments per condition. (B) Colony formation assay of *KDM3A*, *KDM3B* and

*KDM5B* knockout versus wildtype. n=3 independent experiments per condition. (C) Proliferation of *KDM3A*, *KDM3B* and *KDM5B* knockout cells versus wildtype characterized by doubling time. n=3 independent experiments per condition. (D) Cell migration of *KDM3A*, *KDM3B* and *KDM5B* knockout cells versus wildtype by wound healing assay. n=3 independent experiments per condition. Mean ± SD are presented. $p$ value was calculated using unpaired two-tailed Student's t test (C) and (D); ns, not significant; $^{**}p < 0.01$, $^{***}p < 0.001$, $^{****}p < 0.0001$.

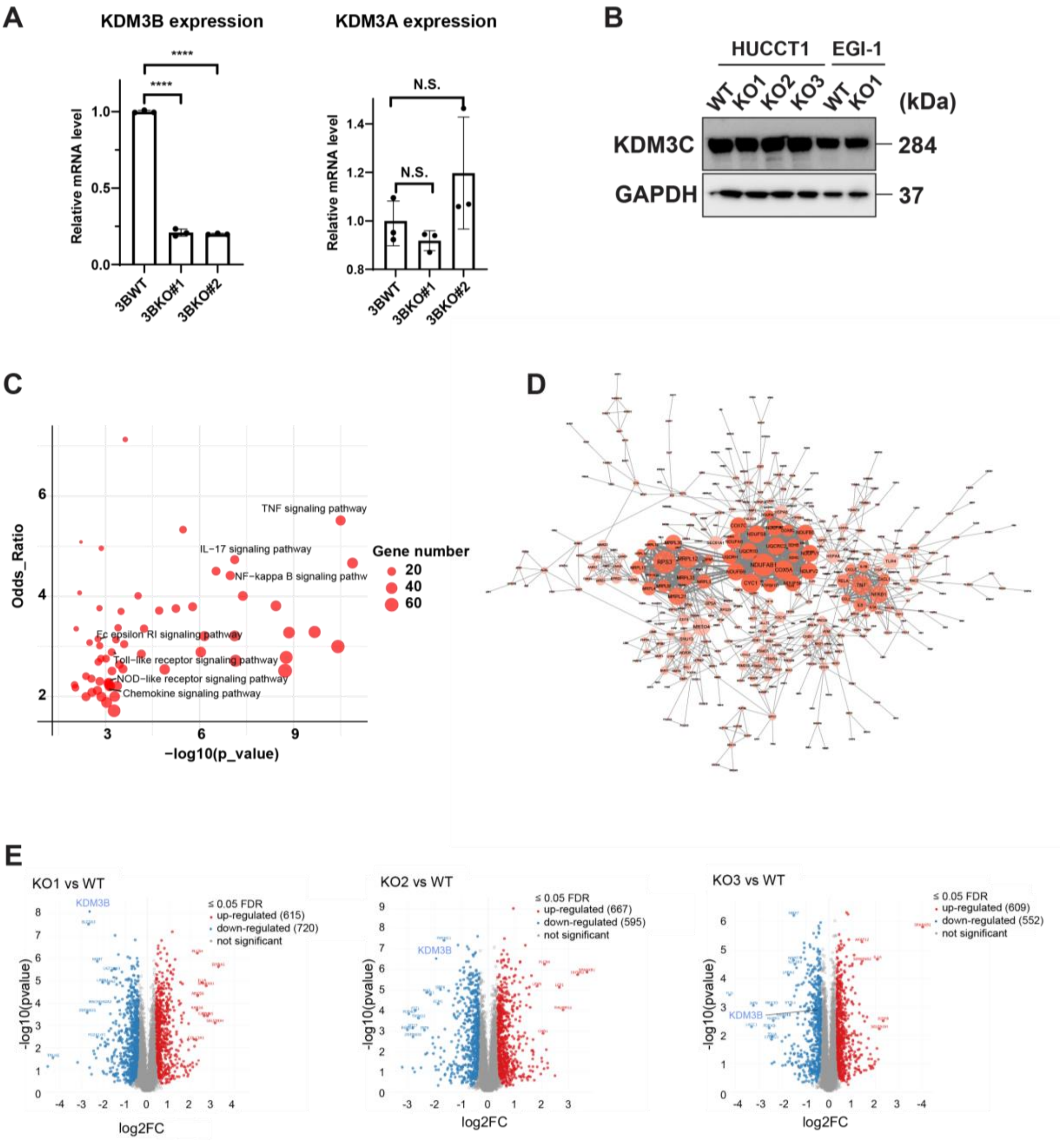

**Figure S3. Transcriptomic and proteomic analysis in *KDM3B* knockout cells, related to Figure 3.** (A) mRNA expressions of *KDM3A* and *KDM3B* in *KDM3B* KO revealed by qRT-PCR. (B) Protein expression of *KDM3C* in wildtype (WT) and *KDM3B* knockout (KO) in HUCCT1 and EGI-1 cell models. (C) KEGG-defined enriched gene sets ($p < 0.05$) among upregulated genes in *KDM3B* knockout cells. Key signaling pathways were highlighted. (D) Protein-protein interaction (PPI) showed the core genes and networks among the up-regulated genes after *KDM3B* knockout. (E) Volcano plot illustrating differentially expressed proteins in

KDM3B knockout (KO1, KO2, and KO3) clones relative to wild-type (WT) cells, as determined by global mass spectrometry-based proteomic analysis.

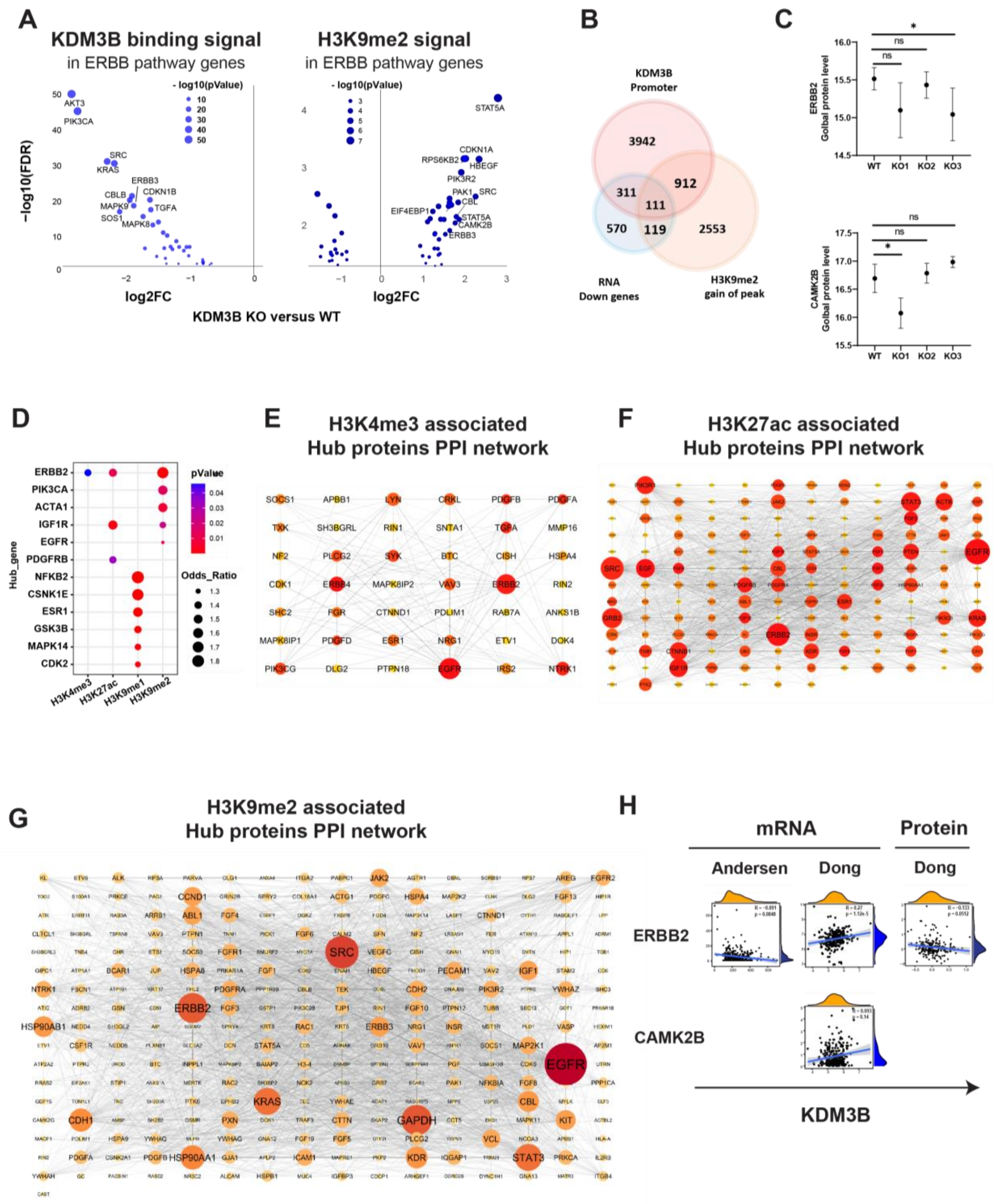

**Figure S4. Extended characterization of CHIP seq profile and patient confirmation, related to Figure 4 and 5.** (A) Volcano plot showed significantly differential genes with KDM3B binding (left panel) and H3K9me2 modification (right panel) in ErbB pathway. (B) Overlap of genes with KDM3B promoter occupancy, transcriptional downregulation, and increased H3K9me2 peaks following KDM3B KO, representing a putative KDM3B target-gene signature. (C) Global protein level of ERBB2 and CAMK2B in wildtype and KDM3B KO models by mass spectrometry. (D) EnrichR-based hub gene analysis of significantly altered peak-associated genes marked by H3K4me3, H3K27ac, H3K9me1, and H3K9me2 ($p < 0.05$). (E) Protein-protein interaction (PPI) network of ERBB2 and proteins associated with differential H3K4me3 occupancy. (F) Protein-protein interaction (PPI) network of hub proteins ERBB2, IGF1R, and PDGFRB and proteins associated with differential H3K27ac occupancy. (G) PPI network of hub genes ERBB2, PIK3CA, ACTA1, IGF1R and EGFR and proteins associated with differential H3K27ac occupancy. (H) Association of *KDM3B* with *ERBB2* and *CAMK2B* at RNA and protein levels in CCA patient cohorts (Andersen, Dong).

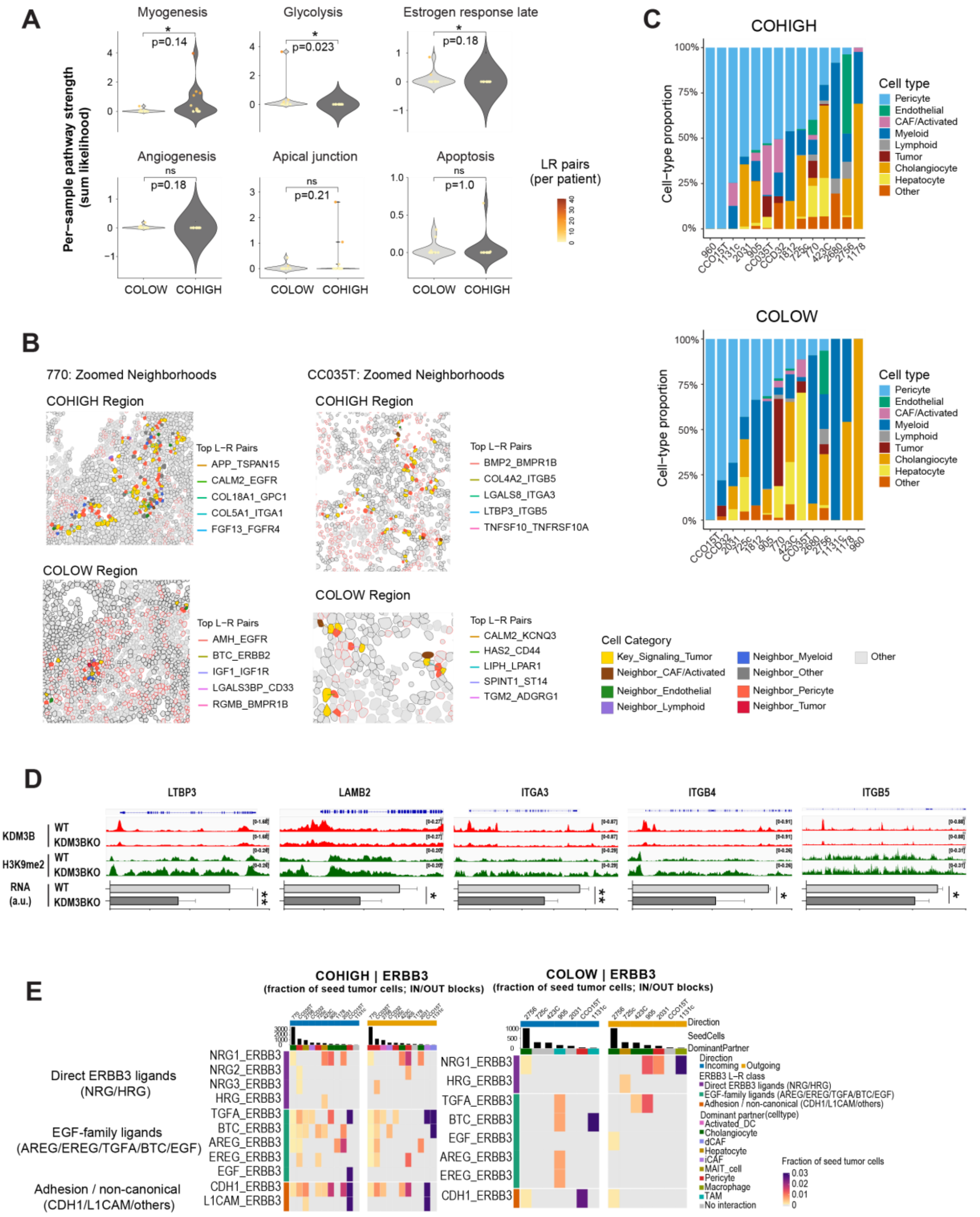

**Figure S5. Spatial transcriptomic analysis of tumor ecosystem associated with KDM3B-ERBB3 expression, related to Figure 6.** (A) Violin plot showed the strength of KDM3B-ErbB3 cohigh and colow in top enriched hallmark gene sets. Paired-Wilcoxon test, n =13. (B)

Representative spatial maps showing top five ligand–receptor (L-R) interactions between tumor cells and adjacent cell types in cohigh and colow regions. (C) Proportions of neighboring cell types in KDM3B–ERBB3 cohigh and colow regions within CCA tissue sections. (D) RNA level and Genome browser view of CHIP-seq signal intensity (KDM3B-, H3K9me2-occupied regions) at the locus of *LTBP3*, *LAMB2*, *ITGA3*, *ITGB4* and *ITGB5* following *KDM3B*$^{-/-}$ in HUCCT1 cells. (E) ERBB3-associated L-R pairs between tumor cells and neighborhood in KDM3B-ERBB3 co-high and co-low regions.

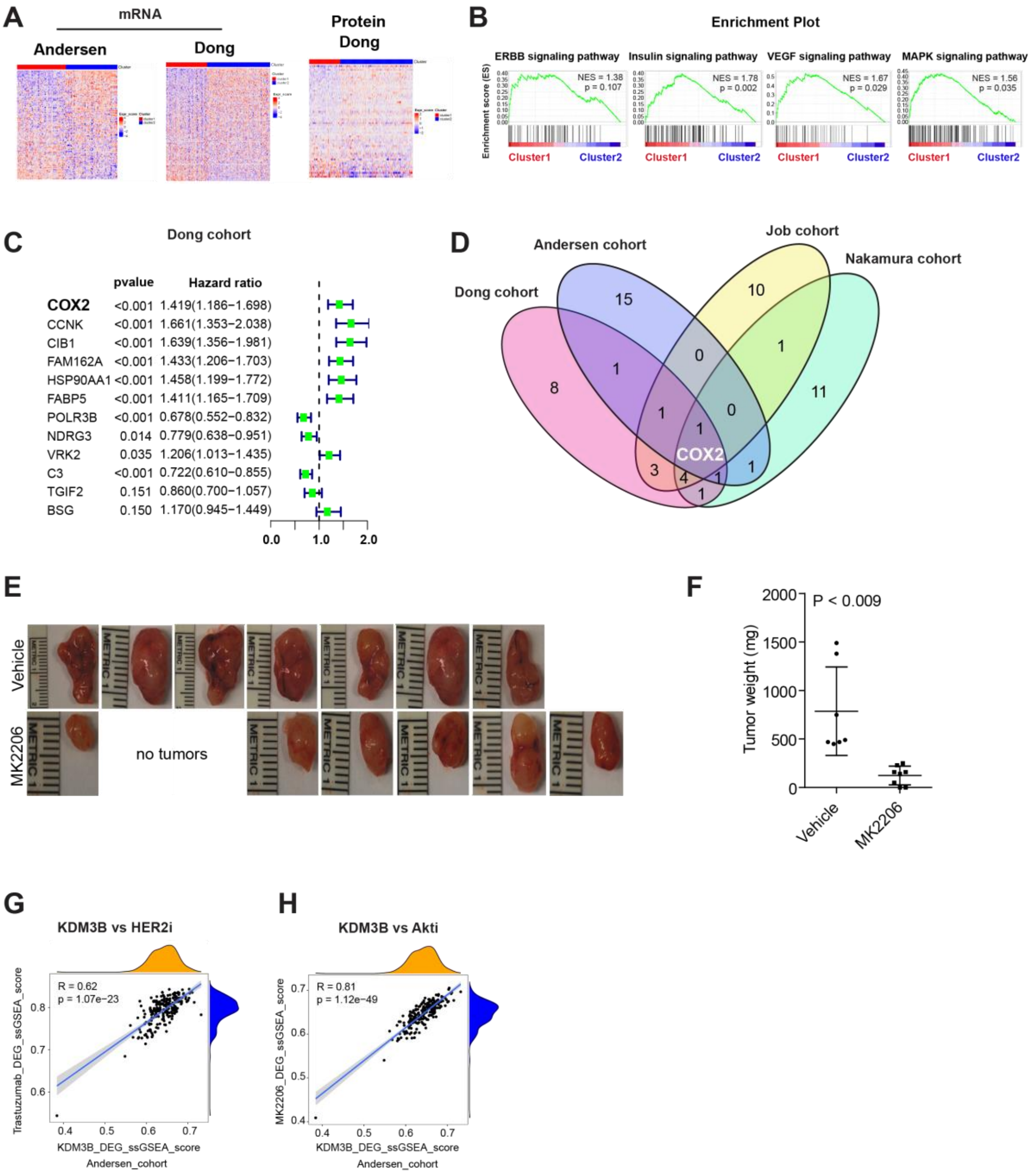

**Figure S6. Targeting the KDM3B-ErbB-Akt axis exhibited anti-tumor effect *in vivo*, related to Figure 7.** (A) Heatmap showing RNA and protein expression of the trastuzumab-response signature across two CCA patient clusters in the Andersen and Dong cohorts. (B) KEGG signaling pathway analysis by gene set enrichment analysis (GSEA) comparing the two clusters in the Dong proteomic cohort. (C) Identification of survival-associated genes within the trastuzumab response signature using univariate Cox analysis in Dong CCA cohort. Forest plot shows risk-associated (HR > 1) and protective (HR < 1) genes. (D) Overlap analysis of survival-associated genes across four CCA cohorts (n = 662) identified COX2 as the consistently associated gene. (E) Xenograft tumor images following treatment with vehicle or the pan-Akt inhibitor MK2206. (F) Tumor weight following treatment with MK2206 or vehicle. (G) Correlation of ssGSEA scores for KDM3B KO DEGs and trastuzumab-responsive DEGs in the Andersen cohort. (H) Correlation of ssGSEA scores for KDM3B KO DEGs and MK2206 (Akti)-responsive DEGs in the Andersen cohort.

### EXPERIMENTAL MODEL AND STUDY PARTICIPANT DETAILS

#### Mouse models

Animals were cared for according to the guidelines established by the National Institute of Health's Guide for the Care and the U.S. Public Health Service policy on Human Care and Use of Laboratory Animals. Athymic NCr-nu/nu mice were between 8-12 weeks of age at the time of transplantation. A total of 1 million HUCCT1 cells were injected into the mouse model for a mimic of cholangiocarcinoma. Cells were resuspended in PBS and mixed with Matrigel at a 1:1 ratio. A final volume of 100ul of the cell suspension was subcutaneously injected into the flanks of the mice. Once tumors reached the indicated volume (40 $mm^3$ for the trastuzumab and combination studies; 200 $mm^3$ for the MK2206 study), mice were randomized into the indicated treatment or vehicle control groups ($n \geq 6$ tumors per group). Trastuzumab was administered intraperitoneally (0.45 mg per injection) twice weekly. Combination-treated mice received celecoxib (30 mg/kg, p.o., twice daily) by oral gavage together with trastuzumab. For AKT inhibition, MK2206 (120 mg/kg, p.o., three times weekly; pan-AKT inhibitor, Merck) was administered by oral gavage. Control mice received vehicle (0.5% methylcellulose). Tumor volume (TV) was measured by caliper measurements and calculated using the formula:

TV = (length × $width^2$)/2. Relative tumor volume (RTV) was determined as the ratio of tumor volume at each time point (week 1-4) to the baseline tumor volume on day 0 ($RTV = TV_t/TV_0$). Mice were observed daily and euthanized when reaching a humane endpoint.

**Cell lines**

The human cholangiocarcinoma cell lines HUCCT-1, EGI-1, TKKK, KMBC, KMCH, CC16, CCSW1 were maintained in Dulbecco's modified Eagle's medium (DMEM; Thermo Fisher Scientific, 31966-047). The human cholangiocarcinoma cell lines, SNU478, SNU1196, SSP25, SNU1079, YSCCC, RBE, SNU308, SNU245, CCLP1 were maintained in RPMI-1640 medium (ThermoFisher Scientific, 72400054). Huh28 was cultivated in RPMI1640 L-glut + 10% heat-inactivated FBS + 1% EpiCGS. Human cholangiocytes (H69), NHC2, NHC3, NHCSS, C324 were cultured in Ham's F-12 medium supplemented with 1 mM L-glutamine, 0.1 mM non-essential amino acids (Thermo Fisher Scientific, 11140050), 1 mM sodium pyruvate (Thermo Fisher Scientific, 11360070), and 10 mM HEPES (Thermo Fisher Scientific, 15630080). Human embryonic kidney 293T cells (HEK293T; CRL-3216, ATCC) were maintained in DMEM. All media were supplemented with 10% fetal bovine serum (FBS; Thermo Fisher Scientific, 10270106), 100 U/mL penicillin, and 100 µg/mL streptomycin (Life Technologies, 15140-122).

### METHOD DETAILS

**Generation of knockout cell lines**

Knockout cell models were generated using the CRISPR-Cas9 system[2]. Guide RNAs (gRNAs) targeting *KDM3A*, *KDM3B*, *KDM5A*, and *KDM5B* were designed immediately downstream of the transcription start site (TSS) using the Integrated DNA Technologies (IDT) CRISPR design tool (https://eu.idtdna.com/site/order/designtool/index/CRISPR_PREDESIGN). The gRNA sequences were cloned into the pLVX-TetOne-Puro vector and transfected into the intrahepatic cholangiocarcinoma (iCCA) cell line HUCCT-1 and the extrahepatic cholangiocarcinoma (eCCA) cell line EGI-1. Successfully transfected GFP-positive cells were enriched by fluorescence-activated cell sorting (FACS). Single-cell clones were isolated by serial dilution and expanded. Genomic DNA was isolated from individual clones, and the targeted regions were amplified by PCR and verified by Sanger sequencing to identify knockout clones. Knockout clones were further confirmed by western blot analysis.

**Epi-drugs sensitivity analysis**

Epi-drug sensitivity was assessed using an in-house drug repositioning screen comprising 525 compounds that are FDA-approved or in late-stage clinical development[3]. A total of 61 epigenetic drugs were selected for evaluation. The screen was performed across eight cholangiocarcinoma (CCA) cell lines (HUCCT-1, EGI-1, SNU478, SUN1079, SNU1196, REB, WITT, and KMCH). Cells were seeded into 384-well plates and treated for 72 hours using a five-fold serial dilution series. Anti-proliferative effects were quantified using the CellTiter-Glo Luminescent Cell Viability Assay (Promega) according to the manufacturer's instructions. Drug responses were summarized using the Drug Sensitivity Score (DSS) metric[4]. DSS values for the compounds were subsequently extracted for downstream evaluation and visualization.

**Expression analysis of CCA cell lines**

Primary human normal cholangiocyte (NHC2, NHC3, NHCSS, C324) and 16 cholangiocarcinoma cell lines (TKKK, Huh28, SNU478, SNU1196, SSP25, HuCCT1, KMBC, SNU1079, CC16, YSCCC, KMCH, CCLP1, RBE, SNU308, SNU245, CCSW1) were collected and maintained in house. Total RNA was isolated using standard procedures, and gene expression profiling was performed by microarray analysis following the protocol described by Jesper et al[5]. Expression of *KDM3A/B* and *KDM5A/B* were withdrawn for visualization.

**Chromatin immunoprecipitation sequencing (ChIP-seq)**

For each sample, $1\times10^7$ cells were crosslinked with 1% formaldehyde (Merck, 252549-100ML) at room temperature for 10 min, and the reaction was quenched with 0.125 M glycine for 2 min. Cells were washed twice with ice-cold PBS with protease inhibitor and either flash-frozen at −80 °C or resuspended in 2 mL freshly prepared SDS lysis buffer (50 mM Tris pH 8.0, 100 mM NaCl, 5 mM EDTA pH 8.0, 0.5% SDS, and protease inhibitor cocktail) and incubated at room temperature for 15 min. Nuclei were pelleted by centrifugation at maximum speed, resuspended in 1 mL sonication buffer (2 volumes SDS lysis buffer + 1 volume ChIP dilution buffer), and transferred into Covaris milliTUBE 1 mL Fiber tubes. Chromatin was sheared to 150-500 bp fragments using a Covaris E220 Evolution sonicator (settings: Peak Power 140, Duty Factor 10%, Cycles per Burst 200; cycles of 60, 120, and 180 s). Lysates were cleared by centrifugation at maximum speed for 15 min, and chromatin concentration in the supernatant was measured by BCA assay. Equal amounts of chromatin were incubated overnight at 4°C with rotation in the presence of specific antibodies (anti-KDM3B abcam ab70797, anti-$H_3K_9me_2$ abcam ab1220, anti-$H_3K_9me_1$ abcam ab9045, anti-$H_3K_4me_3$ CST #9751, anti-$H_3K_{27}ac$ abcam ab4729) together with 2% exogenous mouse chromatin as spike-in control. Two percent of the chromatin was reserved as input. Antibody-chromatin complexes were captured with Dynabeads™ Protein G (Thermo Fisher Scientific, 10004D) for 4 h at 4°C with rotation. Beads were washed sequentially: three times with low-salt buffer (20 mM Tris-HCl pH 8.0, 150 mM NaCl, 2 mM EDTA pH 8.0, 0.1% SDS, 1% Igepal CA-630), twice with RIPA buffer (50 mM Tris-HCl pH 8.0, 150 mM NaCl, 1 mM EDTA pH 8.0, 0.1% SDS, 1% Igepal CA-630, 0.5% sodium deoxycholate), once with low-salt buffer, once with high-salt buffer (20 mM Tris-HCl pH 8.0, 500 mM NaCl, 2 mM EDTA pH 8.0, 0.1% SDS, 1% Igepal CA-630), and twice with low-salt buffer. Immunoprecipitated chromatin and input controls were reverse crosslinked overnight at 65 °C in 100 µL ChIP de-crosslinking buffer (0.1 M $NaHCO_3$, 1% SDS), and DNA was purified using MinElute columns (Qiagen, 28004). For KDM3B ChIP, cells were pretreated with 2 mM disuccinimidyl glutarate (DSG) in PBS for 30 min at room temperature, washed 2-3 times with PBS, and immediately fixed with formaldehyde, followed by subsequent steps described above.

ChIP enrichment efficiency was assessed by qPCR using cycle threshold (Ct) values at a desert region and an enriched region. ChIP was considered successful when the enrichment at the reference target region relative to the desert region exceeded 10-fold. Ct values of the reference enriched region were also used to verify consistency of starting chromatin amounts across samples. Successful ChIP and input DNA were used for library preparation with the NEBNext Ultra II DNA Library Prep Kit for Illumina (New England Biolabs, E7645L) according to the manufacturer's instructions. Libraries were validated using the Agilent High sensitivity DNA kit and the concentrations were measured by Qubit dsDNA HS Assay Kit. Libraries were sequenced on an Illumina NextSeq 500 platform with 42-bp paired-end reads.

**Immunoblotting**

Cells were lysed in RIPA buffer (150 mM NaCl, 1 mM EDTA, 50 mM Tris pH 7.5, 1% sodium deoxycholate, 1% NP-40, 0.1% SDS) supplemented with protease inhibitor cocktail (Complete, Roche Applied Science, 11836153001). Soluble extracts were quantified using the BCA assay. Equal amounts of protein were denatured in 1× SDS loading buffer (250 mM Tris-HCl pH 6.8, 10% SDS, 30% glycerol, 500 mM DTT, 0.05% bromophenol blue), resolved by SDS-PAGE, and transferred onto polyvinylidene difluoride (PVDF) membranes by semi-dry blotting. Membranes were blocked in 3% Bovine Serum Albumin (BSA) for 1 h at room temperature and incubated with primary antibodies overnight at 4°C. After three washes with TBS-T, membranes were incubated with HRP-conjugated secondary antibodies for 1 h at room temperature, washed again, and developed with Immobilon Crescendo Western HRP substrate (Millipore, WBLUR0020).

The following primary antibodies were used: KDM3B (Cell Signaling, #2621, 1:1000), KDM3A (Proteintech, 12835-1-AP, 1:1000), KDM5A (Abcam, ab78322, 1:1000), KDM5B (Merck, HPA027179, 1:1000), GAPDH (Abcam, ab22555, 1:2000), HA (Sigma-Aldrich, H6908-100UL, 1:2000), V5 (Cell Signaling, #13202, 1:1000), KDM3C (Millipore, 09-817, 1:1000), ERBB2 (Cell Signaling, #2165, 1:1000), and ERBB3 (Cell Signaling, #12708, 1:1000). HRP-conjugated secondary antibodies included anti-mouse IgG (Cell Signaling, #7076, 1:5000) and anti-rabbit IgG (Cell Signaling, #7074, 1:5000).

**Reverse-transcriptase quantitative PCR (RT-qPCR)**

Total mRNA was isolated from 1 x $10^5$ to 9 x $10^5$ cells using the RNeasy Kit (Qiagen) according to the manufacturer's instructions. For RT-qPCR, 200 ng of RNA was reverse transcribed and amplified using Luna Universal qPCR Master Mix (New England Biolabs, E3005E). The following primers were used to measure the expression of *KDM3A* and *KDM3B*: *KDM3A* F: TTGAGCCACACAGACAGGTT R: CAGCACCCTGTTGGCAATTC *KDM3B* F: CCCAACATACACCATCAC R: GGGCCCCAAGATCAA. *GAPDH* F: GACAGTCAGCCGCATCTTCT R: TTAAAAGCAGCCCTGGTGAC. Relative expression was calculated using the ΔCt method, with *GAPDH* serving as the internal reference gene.

**Colony formation assay**

A total of 1,500 cells per well were seeded into 6-well plates and cultured for 10-14 days. Cells were gently rinsed with ice-cold PBS, fixed with methanol at -20 ℃ for 10 min, washed three times with PBS and stained with crystal violet solution for 5 min. Plates were rinsed thoroughly with distilled water, air-dried, and imaged. Colony areas were quantified using ImageJ software (NIH, Bethesda, MD, USA).

**Cell proliferation**

A total of 3,000 cells per condition were seeded into 96-well plates and monitored every 4 h for 160 h using the IncuCyte S3 Live-Cell Analysis System (Sartorius). Doubling time was calculated during the exponential growth phase using the formula: $\text{Doubling time} = \frac{\ln 2\ (t2 - t1)}{\ln N2 - lnN1}$, where t represents time and N represents cell confluence.

**Wound healing assay**

A total of 40,000 cells per well were seeded in 100 µL of medium into Incucyte® ImageLock 96-well plates (Essen BioScience, 4379) and incubated overnight to allow attachment. Uniform scratch wounds were generated using the Incucyte® WoundMaker tool, and images were acquired every 40 min using the Incucyte S3 system (Sartorius). Wound width was quantified using Incucyte 2022B Rev2 software, and wound closure was calculated as:

$$\%\ Would\ width = \frac{\text{wound width at 6 h}}{\text{Initial wound width at 0h}} \times 100\%$$

**Transcriptomics**

For RNA sequencing, total RNA was extracted from wild-type (WT) and three *KDM3B* knockout (KO) HUCCT1 clones using the RNeasy Kit (Qiagen) according to the manufacturer's instructions. RNA integrity was assessed with an Agilent 2100 Bioanalyzer (Agilent Technologies), and only samples with RNA integrity number (RIN) >9.5 were used for library preparation. Libraries were generated from 500 ng of high-quality RNA per sample using the Illumina® Stranded Total RNA Prep kit (Illumina, 20040529) following the manufacturer's protocol. Library concentrations were measured using a Qubit Fluorometer (Thermo Fisher Scientific, Q32853), and fragment size distribution was confirmed using a

TapeStation system (Agilent, 5067-5588). Libraries were sequenced on an Illumina NextSeq 550 platform to generate 75-bp paired-end reads.

For microarray analysis, total RNA was isolated from fresh-frozen xenograft tumors treated with trastuzumab, MK2206, or the corresponding vehicle controls. Sample processing was performed as previously described[5]. 200 ng of total RNA was amplified to generate biotin-labeled cRNA according to the manufacturer's protocol. A total of 750 ng biotinylated cRNA was hybridized to HumanRef-8 v2 BeadChips (Illumina, San Diego, CA, USA) at 58°C for 18 h. Arrays were subsequently washed, stained with Cy3-streptavidin, and scanned using an Illumina iScan System. Raw fluorescence intensities were extracted using Illumina iScan Control Software.

**Immunofluorescence**

Formalin-fixed paraffin-embedded (FFPE) tumor tissues were obtained from patients with good prognosis and poor prognosis defined by the CCA signature[5]. FFPE blocks were cut into sections, mounted on glass slides, deparaffinized, and subjected to antigen. All sections were processed under standardized conditions to minimize batch variability. After blocking, sections were incubated with primary antibodies against COX2 (Novus Biological, # NB100-689) and additional tumor marker CK19 (Proteintech, 10712-1-AP) as indicated. Following thorough washes, slides were incubated with the Alexa Fluor-conjugated secondary antibodies for 1 h at room temperature. Nuclei were counterstained with DAPI, and slides were mounted with antifade medium.

Images were acquired using a confocal under identical laser power and exposure settings across experimental groups. Representative fields of view were collected in a blinded manner. Fluorescence signal intensity was evaluated qualitatively by visual inspection of representative fields by observers blinded to sample identity.

**Global proteomics and Phosphoproteomics**

***Sample preparation:*** Cell pellets were lysed by the addition of preheated (95 °C) lysis buffer (5% SDS, 100 mM Tris, pH 8.5) and homogenized with a BeatBox homogenizer (PreOmics) for 5 min at the high setting. Samples were incubated at 95 °C for 10 min and lysed with a second round of BeatBox for 10 min at standard setting, followed by incubation at 95 °C for 10 min. Protein concentrations were determined using the BCA assay (Pierce). Samples were reduced with 5 mM Tcep (final concentration) for 15 min at 55 °C, and alkylated with 20 mM CAA (final conc.) for 30 min at room temperature. 233 µL acetonitrile (ACN), and 25 µL of pre-equilibrated (70% ACN) MagReSyn Hydroxyl beads (Resyn Biosciences) were added to 100 µL protein extract (100 µg protein), and the samples were mixed and settled for 10 min at room temperature. Beads were washed three times with 1 mL 95% ACN, twice with 1 mL 70% EtOH, and digested overnight at 37 °C with 100 µL digestion buffer (50 mM TEAB) containing Trypsin and Lys-C in a 1:50 (enzyme: protein) ratio. Peptides were acidified with 1% (final conc.) TFA and desalted using EasyPep peptide clean-up plates (Thermo Fisher) following the manufacturer's instructions. Eluates were dried using a vacuum centrifuge and resuspended in 20 µL of 0.1% TFA containing 0.01% DDM for phosphopeptide enrichment.

Phosphorylated peptides were enriched using Fe(III)-NTA cartridges (Agilent Technologies) in an automated manner on the AssayMAP Bravo Platform (Agilent Technologies), following the standard protocol. Briefly, cartridges were primed with 100 µL of priming buffer (100% acetonitrile, 0.1% TFA) and equilibrated with 50 µL of loading buffer (80% acetonitrile, 0.1% TFA). After sample loading, cartridges were washed with the loading buffer while syringes were rinsed with the priming buffer. Phosphopeptides were eluted using 20 µL of 500 mM ammonium dihydrogen phosphate ($NH_4H_2PO_4$) containing 0.01% DDM. Both the enriched phosphopeptides and flowthroughs were dried by vacuum centrifugation and resuspended in buffer A* (0.1 M TFA, 2% ACN).

***LC-MS analysis:*** Peptides were separated on a C18 PepMap trapping column, 300µm inner diameter (Thermo Scientific) using Vanquish Neo UHPLC system (Thermo Scientific), followed by separation on an Aurora Ultimate 25×75 C18 column (IonOpticks), Peptide separation was performed using a 60 min stepped gradient of 2-17% solvent B (0.1% formic acid in acetonitrile) for 33 min, 17-25% solvent B for 11 min, 25-35% solvent B for 6 min, using a constant flow rate of 400 nL/min. Column temperature was controlled at 50 °C. Upon elution, peptides were injected into a Tribrid Ascend mass spectrometer (Thermo Scientific). Spray Voltage was set to 2200 (V). Data was acquired in data independent mode with the Orbitrap MS resolution 60.000 for full scan range 400-900 m/z for the global proteome and 450-1150 m/z range for the phospho proteome, AGC target was 250% and maximum injection time 123 ms. For global proteome data, 41 DIA scans with 12 Th width and 1 Th overlap spanning a mass range of 400-900 m/z were acquired at 15,000 MS resolution, AGC target 1000% and maximum injection time of 27 ms. CID fragmentation normalized collision energy (NCE) was set to 30%. For phosphorylation, 55 DIA scans a mass range of 472-1143 m/z were acquired with the same settings as described above.

**Patient characterization and omic datasets**

The Andersen cohort comprised 210 tumors (T) and 143 matched surrounding liver (SL) tissues, as previously described[3,5]. The spatial transcriptomics include tumor tissue from 14 CCA patients[6]. Ethics approval and written informed consent were obtained in accordance with local regulations, and the study was approved by institutional review boards at the contributing institutions. Whole-transcriptome profiling was performed on the Illumina platform, and data are available under accession number GSE265664[3,5]. Publicly available datasets were included for comparative analyses: the Dong cohort (255 iCCA)[7], the Job cohort (E-MTAB-6389[8], 78 iCCA and 31 surrounding liver tissues), TCGA (29 iCCA and 6 eCCA)[9], TIGER (91 tumors and 92 matched normal tissues)[10] and the Montal cohort (182 eCCA and 38 surrounding bile duct tissues)[11], Single-cell RNA-seq datasets were obtained from GSE125449[12], GSE138709[13], GSE146409[14], GSE151530[15] and GSE136103[16].

**RNA-seq analysis**

Raw RNA-sequencing data were assessed for quality using FastQC (v0.11.9). Sequencing adapters were removed using trimmomatic. Cleaned reads were aligned to the human reference genome (GRCh38) and quantified using the Rsubread package (v2.18.0). Differential gene expression analysis was performed using DESeq2 (v1.42.1). Functional enrichment analyses, including KEGG pathway clustering, and PPI hub protein analysis, were conducted using Enrichr (https://maayanlab.cloud/Enrichr/)[17].

**CHIP-seq analysis**

ChIP-seq data were processed using the nf-core/chipseq pipeline (v2.0.0)[18]. Raw reads were aligned to the human reference genome (GRCh38) with Bowtie2 (v2.4.4), and duplicate reads were marked and removed using Picard (v2.27.4). Peak calling was performed with MACS2 (v2.2.7.1). For KDM3B, H3K4me3, and H3K27ac ChIP, sharp peaks were identified using the parameters --narrow_peak --macs_fdr 0.05; for H3K9me1 and H3K9me2 ChIP, broad peaks were identified using --broad_cutoff 0.1. Genome-wide coverage tracks were generated with BEDTools genomecov (v2.30.0) to produce BedGraph files, which were converted to BigWig format using BedGraphToBigWig (UCSC, v377). BigWig signals were normalized to one million reads for cross-sample comparison.

Data visualization was performed with Integrative Genomics Viewer (IGV, v2.15.4), and heatmaps were generated using deepTools (v3.5.1). Differential peak analysis between WT and *KDM3B* KO samples was performed with DESeq2 (v1.42.1). Genomic feature annotation and motif discovery were carried out using the annotatePeak.pl script from HOMER (v4.11). Motif analysis of KDM3B binding was performed using all peaks from WT samples with peaks from *KDM3B* KO samples as background.

**Proteomic and Phosphoproteomic analysis**

***Data Processing***: MS files for the global proteome were processed using Spectronaut version 19.9.250422 (Biognosys) in direct DIA search mode with the default "BGS Default Workflow". Briefly, UniprotKB UP0000005640 (Taxa ID: 9606) Homo sapiens FASTA database was used. Carbaminomethylation of cysteine was set as a fixed modification, whereas the oxidation of methionine, the acetylation of the protein N-terminus were set as variable modifications. For phosphoproteomic data, the direct DIA search mode with the default "BGS Phospho PTM Workflow" was used, with phosphorylation of tyrosine, serine, and threonine set as variable modifications. For both searches, the maximum number of missed cleavages was 2, and the minimum peptide length was 7 amino acids. The false discovery rate for PSM, peptide, and protein groups was set to 0.01. The PTM localization was activated, and the site confidence score cutoff was set to 0.75 for phosphoproteomics data.

***Data analysis***: All statistical analysis was performed using in-house developed python code (Proteomelit, version 1.1.0), based on the automated analysis pipeline of the Clinical Knowledge Graph[19]. For phosphoproteomic data, precursors were converted to modified peptides by grouping unique modified sequences in each sample and summing the precursor intensities. Intensity values were log2-transformed, and features with less than 70% of valid values in at least one group were removed. For the phosphoproteomes dataset, statistical and enrichment analyses were performed on phospho-modified peptides only. Remaining missing values were replaced by mixed imputation, where kNN and MinProb (width=0.2 and shift=1.8) methods are used for values missing at random (MAR) and values missing not at random (MNAR), respectively[20]. MAR is defined when a minimum of 60% of the samples within a given group have an existing value. Once processed, modified peptides were normalized to the protein abundance in the non-enriched global proteome. Differentially expressed features were identified by unpaired t-tests. p-values were corrected for multiple hypothesis testing with permutation-based False Discovery Rate (FDR) correction. False Discovery Rate (FDR) threshold was set to 0.01, Fold-change (FC) 1, s0 1, and 250 permutations.

Functional enrichment analysis of significant features was performed by applying Fisher's exact test and Benjamini-Hochberg correction. Kyoto Encyclopedia of Genes and Genomes (KEGG) were used for the enrichment testing. The analysis was performed at the protein level for both the global proteome and phosphoproteome datasets. Up- and down-regulated features were used as foreground, and the remaining features in each dataset were considered background. For the phosphoproteome dataset, linear motif enrichments were performed using the same strategy. Annotation data were obtained from PhosphoSitePlus[21] and Phosida[22], and phosphorylated protein sites mapped to significantly regulated modified peptides were used as foreground. Enrichment bubble plots display only significantly enriched terms (adjusted p-value $< 0.05$) with at least 2 proteins of the foreground associated with the term. Kinase activity inference was performed using Fisher's enrichment analysis[23,24]. Changes in phosphorylation levels of known kinase substrates were used to infer kinase activity. The inferred kinase activity changes were visualized on a kinome tree using Coral[25].

**Single-cell RNA-seq (scRNA-seq) analysis**: Single-cell transcriptomic data were obtained from 27 CCA tumors (T) and matched adjacent normal tissues (SL), comprising a total of 44,761 cells (GSE151530[15], GSE138709[13], GSE146409[14], GSE125449[12]). Seurat4 pipeline was used for processing datasets. Data processing and downstream analysis were performed using the Seurat R package (v4.0.0). Individual datasets were merged, and normalization and scaling were performed with SCTransform, regressing out mitochondrial RNA percentage and cell cycle effects. Known marker genes were used to annotate clusters to cell types. The 3,000 most variable genes were used for principal component analysis (PCA, 35 dimensions), and clustering was performed at a resolution of 0.7. Cell clusters were annotated based on canonical marker gene expression. The average expression levels and the percentage of cells expressing *KDM3A, KDM3B, KDM5A,* and *KDM5B* were compared across clusters and between tissue types.

Cholangiocyte clusters were subset and re-integrated using the Seurat anchor-based integration workflow. PCA (20 components) with 3,000 variable features was applied to generate integration anchors across datasets, followed by dimensionality reduction and clustering on the integrated assay (35 dimensions, resolution = 0.8). Cell-cell communication was performed using the CellChat[26] package. Within the tumor subset, cells were stratified into high and low groups based on the top 25% and bottom 25% of KDM3B target gene expression levels, respectively. Differential signaling pathways were identified by comparing the overall information flow between these two groups. The secreted signaling database was used to assess differential interactions between high and low KDM3B-expressing tumors and their surrounding cellular environments. Correlations between gene expressions were evaluated by linear regression in tumor cells.

**Gene dependency analysis**

Gene dependency data were obtained from the DepMap Portal (https://depmap.org/portal/). Publicly available CRISPR-Cas9-based cell viability screening data from the DepMap 25Q3 were downloaded. Biliary tract cancer cell lines were selected by filtering samples annotated under the biliary tract lineage. CRISPR gene effect scores were extracted for a curated set of 263 epigenetic regulators. Gene effect scores, as defined by DepMap, represent the relative impact of gene knockout on cell viability, with more negative scores indicating increased dependency. The resulting gene dependency matrix was used for visualization via heatmap.

**Copy number alteration analysis**

Copy number alteration data were obtained from the DepMap Portal (https://depmap.org/portal/), using the DepMap 25Q3 OmicsCNGeneWGS dataset. Biliary tract cancer cell lines were selected by filtering samples annotated under the biliary tract lineage. Gene-level copy number values of *KDM3A/B* and *KDM5A/B* were extracted for visualization.

**Protein-protein interaction (PPI) network analysis**

Differentially expressed genes (upregulated and downregulated) following KDM3B knockout were submitted separately to the STRING database (https://string-db.org/) using the highest confidence threshold (score $\geqslant$ 0.900) to construct PPI networks. Networks were imported into Cytoscape (v3.10.2), and interaction scores were analyzed using the CytoNCA plugin. Hub genes and subnetworks were identified and ranked with the Maximal Clique Centrality (MCC) algorithm implemented in the CytoHubba plugin. PPI networks were visualized in Cytoscape, with node size scaled according to interaction degree derived from the calculated scores.

**Gene Set Enrichment Analysis (GSEA)**

GSEA was performed using the GSEA software (v4.3.3, Broad Institute) on ranked bulk transcriptomic profiles from the Andersen and Dong cohorts and on the protein expression dataset from the Dong CCA cohort[7]. Genes were ranked using the Signal2Noise metric between conditions, and enrichment scores were calculated with 1,000 phenotype permutations. Gene sets were obtained from KEGG (v2025.1), and pathways with $p < 0.05$ were considered significant.

**Gene and gene set association in patient dataset**

For bulk transcriptomic and proteomic data from CCA cohorts and transcriptomic data from TCGA, normality was assessed using the Shapiro-Wilk test. Pearson correlation was applied to normally distributed data, whereas Spearman correlation was used for non-normally distributed data. Associations between gene sets were evaluated using single-sample gene set enrichment analysis (ssGSEA) implemented in the GSVA package (v2.3.1). Genes with variance < 0.5 were filtered out. ssGSEA scores were computed for each gene set in individual patients, and pairwise correlations between signatures were assessed using Spearman's rank correlation coefficient.

**Visium HD spatial transcriptomic data processing**

Spatial transcriptomic data were generated in-house with Fresh-frozen iCCA tissues (n = 14), using the 10x Genomics Visium HD platform with CytAssist, as previously reported[6]. Raw FASTQ files were processed using Space Ranger (v4.0.1) and aligned to the human GRCh38-2024-A reference genome with the Visium Human Transcriptome Probe Set v2.1.0. CytAssist-based image registration was performed using 20× brightfield H&E images for spatial alignment.

Single-cell segmentation and spatial gene expression assignment were performed using the ENACT (Enhanced Nuclear and Cytoplasmic Assignment for Transcriptomics) pipeline. Tissue segmentation was conducted using StarDist ("2D_versatile_he" model) on H&E images, followed by cytoplasmic expansion for whole-cell boundary estimation. Gene expression from 2 μm × 2 μm Visium HD bins was assigned to segmented cells using area-weighted allocation based on bin-cell overlap. Highly variable genes were selected (top 25,000), and principal component analysis was performed with components retained to explain ⩾90% cumulative variance (maximum 250 PCs). Clustering was performed using the Leiden algorithm, with resolution selected based on clustering stability across bootstrap resampling. All stochastic steps used a fixed random seed.

Cells were filtered based on total UMI counts, detected genes, and mitochondrial gene percentage. Minimum thresholds were set at ⩾100 genes and ⩾100 UMIs per cell, with an upper gene threshold of 10,000 to exclude potential doublets. Local spatial outlier detection was performed using neighborhood-based modified z-scores. Potential doublets were identified using statistical, quantile-based, and density-based (DBSCAN) approaches. Only cells passing all filters were retained.

Cell-cell spatial adjacency matrices were constructed using Euclidean distances between cell centroids. Neighbor relationships were defined using an adaptive distance threshold derived from nearest-neighbor statistics per sample.

Large-scale chromosomal copy number variation (CNV) was inferred using inferCNV with patient-matched normal epithelial cells within each section as the diploid reference. Genes were ordered by chromosomal position (GRCh38), low-expression genes were filtered, and the i6 HMM model was applied. A CNV burden score was computed per epithelial cell, and cells exceeding predefined burden thresholds were classified as malignant.

Following quality control filtering and CNV-based malignancy classification, all cells were annotated using a curated set of liver cancer-specific marker genes derived from published liver cancer single-cell atlas data. The marker compendium encompassed major lymphoid, myeloid, parenchymal, and stromal populations. The overall annotation framework was adapted from Ruidong et al[27].

Cell type assignment followed a three-tier hierarchical strategy prioritizing liver-specific parenchymal and structural cells over immune populations. Cell type scores were calculated using a marker specificity-weighted approach where marker weights were inversely proportional to their frequency across all cell type signatures.

##### ***KDM3B–ERBB3*** **co-distribution analysis**

The ERBB program was defined as the union of unique gene symbols across eleven MSigDB gene sets selected for ERBB/ERBB3 relevance: Biocarta_ERBB3_pathway, GOBP_ERBB3_signaling_pathway, GOBP_ERBB_signaling_pathway, GOBP_negative_regulation_of_ERBB_signaling_pathway, GOBP_positive_regulation_of_ERBB_signaling_pathway, GOBP_regulation_of_ERBB_signaling_pathway, GOMF_ERBB_3_class_receptor_binding, KEGG_ERBB_signaling_pathway, PID_ERBB2_ERBB3_pathway, Reactome_downregulation_of_ERBB2_ERBB3_signaling, WP_ERBB_signaling, yielding a union of 218 genes as ERBB3 program (Table S5C). The KDM3B activity was defined by the KDM3B-target gene signature as described in the main text (Figure S5B and Table S4B).

For each specimen, CNV-classified tumor cells were stratified using within-sample quartiles of QC-adjusted KDM3B (K_adj) and ERBB3 (E_adj) scores: cells ⩾75th percentile for both scores were defined as COHIGH, and cells ⩽25th percentile for both as COLOW, with thresholds computed independently per sample to avoid cross-sample bias.

To assess whether elevated KDM3B activity co-occurs with ERBB3 program activation beyond chance, tumor cells were stratified per sample using quantile-based gating of QC-adjusted scores. Coupling was evaluated using a 2×2 contingency framework comparing the observed high–high co-occurrence rate to that expected under independence, with fold-enrichment defined as p_obs/p_exp. Statistical significance was determined by one-sided Fisher's exact test. To exclude effects of quantile thresholding or residual QC confounding, a continuous association analysis was performed. Ranked KDM3B and ERBB3 scores were residualized against ranked QC covariates, and partial Spearman correlation (ρ) was computed. Samples were required to meet ρ ⩾ 0.20 and ⩾15 cells in both COHIGH and COLOW groups for downstream analysis.

As a minimal-assumption complement to signature scoring, tumor cells were stratified by single-gene KDM3B and ERBB3 expression. Log1p-transformed per-cell values were QC-adjusted using the same rank-based residualization procedure. Within each sample, cells were classified as HIGH (≥75th percentile) or LOW (≤25th percentile), and coupling and differential analyses were performed as for the program-based approach.

**Spatial ligand-receptor and cell-cell communication analysis**

***Ligand-receptor interaction inference between COHIGH and COLOW tumor seed populations:*** Spatially resolved ligand–receptor (LR) interactions were inferred using SCOTIA with a curated human LR reference from CellChatDB v2. SCOTIA constructs adaptive spatial neighborhoods based on local cell density and scores interactions using an optimal transport framework integrating ligand/receptor expression, spatial proximity, and cell-type compatibility. For each cell, outgoing (n_interactions_sent) and incoming (n_interactions_received) interactions were recorded. Interactions were aggregated by source–target cell-type pairs to quantify directional communication axes, summarized by the number of distinct LR pairs and their total and mean interaction likelihood. Neighborhood composition for each central cell was defined by enumerating interaction-linked neighbor cell types, with neighbor counts computed separately for outgoing and incoming edges and then combined into a per-cell neighbor vector.

To compare communication between COHIGH and COLOW tumor cells, analyses were restricted to SCOTIA edges incident on tumor seed cells and performed in a direction-aware manner (OUTGOING: seed as source; INCOMING: seed as target). Within each sample and direction, LR pairs were summarized by the number of unique tumor seed cells involved (primary support metric), supporting edge count, and summed interaction score. LR pairs were globally ranked by pooled seed-cell support, with summed score as a secondary criterion, and a fixed top-40 set per direction was selected for standardized cross-sample comparison.

Using the fixed top-40 LR universe, four cross-sample summaries (COHIGH/COLOW × OUTGOING/INCOMING) were generated. For each LR pair, summaries report unique tumor seed cells stratified by sample, partner cell-type composition (targets for OUTGOING; sources for INCOMING), and per-sample totals including overall tumor seed-cell support and LR-pair diversity.

***Statistical comparison of L-R composition between COHIGH and COLOW:*** To evaluate shifts in ligand-receptor (LR) interaction networks within the selected top-40 LR pairs for either COHIGH and COLOW, a compositional analysis was performed utilizing Cramér's V to quantify effect size. For each directed cell-type pair and enriched biological pathway, interaction frequencies (edge counts) of specific LR pairs were aggregated across the COHIGH and COLOW conditions to construct contingency tables. The statistical significance of compositional differences between these two states was assessed using a Monte Carlo-simulated Chi-square test (B = 4000). To determine the magnitude of these usage shifts independent of total edge volume, Cramér's V was calculated as V = sqrt (x^2 / n), where n is the total number of observations. This yields a standardized metric of network reconfiguration, allowing for the robust identification of state-dependent LR communication changes after applying Benjamini-Hochberg false discovery rate (FDR) correction.

***Pathway enrichment of compartment-level communication arrows and visualization:*** Signaling programs underlying each compartment-to-compartment arrow were assessed by over-representation analysis (ORA) of contributing ligand and receptor genes against MSigDB Hallmark gene sets. The background comprised all unique LR genes detected across SCOTIA edges. Enrichment was tested using one-sided hypergeometric tests (minimum overlap $\geq 2$

genes; minimum restricted gene set size ⩾ 5), with Benjamini–Hochberg correction. Enrichment scores were defined as $-\log_{10}$(adjusted p-value), and each arrow was annotated with its top three pathways; arrows without significant enrichment were labeled accordingly.

For each sample, tumor seed cells defined central neighborhoods (central cell plus interaction-linked neighbors), and only SCOTIA edges with both source and target within the same neighborhood were retained. Edges were aggregated into directional arrows between nine broad cell-type compartments. For each arrow, edge counts and summed interaction likelihoods were accumulated across LR pairs and samples. Arrow thickness was calculated on a shared COLOW–COHIGH scale as the mean of rescaled log1p edge counts and summed likelihood, binned into shared quantiles and mapped to a fixed width range (0.55–4.80 mm).

To evaluate pathway-level interaction strengths between the COHIGH and COLOW states within the selected top-40 LR pairs for either COHIGH and COLOW, a weighted likelihood analysis was performed at the sample level. For each biological pathway, the raw interaction strength was quantified by aggregating the sum of ligand-receptor likelihoods across all directed cell-type edges assigned to that pathway within a given sample. Statistical differences in pathway interaction strengths between the two conditions were assessed using a Wilcoxon test, applying a paired signed-rank test when sufficient matched samples were available ($n \geq 3$) or an unpaired rank-sum test otherwise. Effect sizes were evaluated using mean differences and log2 fold changes, and p-values were adjusted using the Benjamini-Hochberg false discovery rate (FDR) correction to robustly identify condition-specific shifts in overall pathway communication.

***Identification and visualization of spatial seed neighborhood hotspots:*** Hotspot detection was performed per sample and seed set to identify spatially dense tumor niches for visualization. KDM hotspots were identified independently for each gene using tumor-cell-only, multiscale fixed-radius neighbourhood analysis of expression z-scores. Hotspot seeds required elevated local mean expression, detection frequency, and sufficient neighbourhood size. Connected regions were merged across radii when their overlap coefficient was ≥0.60. For each tumor seed cell, a seed-central neighborhood was defined as the central tumor cell plus all SCOTIA-linked neighbors. For each center, neighborhood pericyte fraction, total neighborhood size, and spatial coordinates were computed.

Candidate centers were filtered to those with pericyte fractions between 0.30 and 0.50, consistent with perivascular niches; thresholds were relaxed if fewer than five candidates met this criterion. Spatial clustering was performed using a graph-based connected components approach across a radius grid defined as 0.9–4.0× the median nearest-neighbor distance. For each radius, the largest component was evaluated using a density enrichment metric (Poisson-expected versus observed mean nearest-neighbor distance), and the final hotspot was selected as the largest component with enrichment > 1, or otherwise the radius with maximal enrichment.

Within the selected hotspot, SCOTIA edges were restricted to those whose source or target fell within hotspot neighborhoods. LR pairs were ranked by summed interaction likelihood, and the top five were selected for visualization.

Spatial rendering was performed in R using the sf package. Segmentation-derived cell polygons were overlaid with seed labels and SCOTIA edges, and zoomed dual-panel maps (COHIGH and COLOW) were generated for direct comparison.

***Transcriptional characterization of spatially communicating tumor cells:*** Tumor cells participating in COHIGH and COLOW spatial communication programs were extracted from SCOTIA edge tables in a direction-aware manner. The incoming and outgoing were combined into a union seed set. Cell identifiers were intersected with the CNV-classified malignant compartment per sample to confirm tumor identity.

Per-sample log1p-normalized expression matrices were generated from confirmed malignant COHIGH and COLOW tumor cells. Genes with finite variance were retained, and sample identity was included as a blocking factor. Differential expression was performed using limma with empirical Bayes moderation. When applicable, within-sample inter-cell correlation was estimated using duplicate Correlation. Z-scored log1p library size and detected gene count were included as covariates. The primary contrast tested COHIGH minus COLOW, with Benjamini–Hochberg correction. Analyses required ⩾25 cells per group.

#### QUANTIFICATION AND STATISTICAL ANALYSIS

All bar graphs are presented as mean ± standard deviation (SD). Statistical analyses were performed using GraphPad Prism (v9.3.1) or R (v4.3.3). Comparisons between two groups were assessed using Student's t-test for normally distributed data or Mann-Whitney U test for nonparametric data. For matched-pair analyses, the Wilcoxon signed-rank test was applied. One-way ANOVA was used for comparisons among three or more groups. Survival analyses were performed using Kaplan-Meier estimates, and group differences were assessed with the log-rank test.

Statistical significance was defined as $p < 0.05$, with thresholds indicated as $p < 0.05$ (*), $p < 0.01$ (**), $p < 0.001$ (***), and $p < 0.0001$ (****); non-significant results were reported as n.s.

Multiple testing correction was performed using the Benjamini-Hochberg procedure to control the false discovery rate (FDR). The specific statistical tests used are indicated in the figure legends or results section.

Hierarchical clustering was applied to classify patients based on gene expression signatures. Least absolute shrinkage and selection operator (LASSO) regression was used to identify the most significant genes among trastuzumab-associated signatures linked to survival, and prognostic significance of candidate genes was further evaluated by univariate survival analysis. Heatmaps and correlation plots were generated using the pheatmap and ggplot2 packages in R.

#### Key resources table

| REAGENT or RESOURCE | SOURCE | IDENTIFIER |
|---|---|---|
| **Antibodies** | | |
| Rabbit anti-KDM3A | Proteintech | Cat#12835-1-AP |
| Rabbit anti-KDM3B (WB) | Cell Signaling Technology | Cat#2621 |
| Rabbit anti-KDM3B (CHIP) | Abcam | Cat#ab70797 |
| Rabbit anti-KDM3C | Millipore | Cat#09-817 |
| Mouse anti-KDM5A | Abcam | Cat#ab78322 |
| Rabbit anti-KDM5B | Merck | Cat#HPA027179 |
| Rabbit anti-H3K9me1 | Abcam | Cat#ab9045 |
| Rabbit anti-H3K9me2 | Abcam | Cat#ab1220 |
| Rabbit anti-H3K4me3 | Cell Signaling Technology | Cat#9751 |
| Rabbit anti-H3K27ac | Abcam | Cat#ab4729 |
| Rabbit anti-ERBB2 | Cell Signaling Technology | Cat#2165 |
| Rabbit anti-ERBB3 | Cell Signaling Technology | Cat#12708 |
| Rabbit anti-COX2 | Novus Biological | Cat#NB100-689 |
| Rabbit anti-CK19 | Proteintech | Cat#10712-1-AP |
| Rabbit anti-pAkt(Ser473) | Cell Signaling Technology | Cat#3787 |
| Rabbit anti-GAPDH | Abcam | Cat#ab22555 |
| anti-mouse IgG, HRP-linked | Cell Signaling Technology | Cat#7076 |
| anti-rabbit IgG, HRP-linked | Cell Signaling Technology | Cat#7074 |
| **Biological samples** | | |
| Cholangiocarcinoma patient tissue | Laboratory of Experimental Carcinogenesis, National Cancer Institute and Mayo Clinic | |
| **Chemicals, peptides, and recombinant proteins** | | |
| Crystal violet | Sigma-Aldrich | Cat#C0775 |
| Formaldehyde | Sigma-Aldrich | Cat#252549 |
| Polybrene | Sigma-Aldrich | Cat#H9268-10G |
| Trastuzumab (Herceptin®) | Genentech | N/A |
| MK2206 | MedChemExpress | Cat#HY-10358 |
| cOmplete™, EDTA-free Protease Inhibitor Cocktail | Roche | Cat#05056489001 |

| | | |
|---|---|---|
| Dynabeads™ Protein G | Thermo Fisher | Cat#10004D |
| DreamTaq Green PCR Master Mix | Thermo Fisher | Cat#K1081 |
| HRP substrate | Millipore | Cat#WBLUR0500 |
| D5000 ScreenTape Reagents | Agilent | Cat#5067-5589 |
| D5000 ScreenTape | Agilent | Cat#5067-5588 |
| D5000 Ladder | Agilent | Cat#5067-5590 |
| RNA ScreenTape | Agilent Technologies | Cat#5067-5576 |
| Igepal CA-630 | Sigma-Aldrich | Cat#I8896 |
| Disuccinimidyl glutarate (DSG) | Thermo Fisher | Cat#20593 |
| **Critical commercial assays** | | |
| Illumina® Stranded Total RNA Prep, Ligation with Ribo-Zero Plus | Illumina | |
| NEBNext® Ultra™ II DNA Library Prep Kit | Bionordika (NEB) | Cat#E7103 |
| Qubit™ RNA BR Assay Kit | Invitrogen | Cat#110210 |
| Qubit® dsDNA HS Assay Kit | Invitrogen | Cat#Q32854 |
| QIAquick PCR Purification Kit | Qiagen | Cat#28104 |
| Luna® Universal One-Step RT-qPCR Kit | New England Biolabs | Cat#E3005E |
| RNeasy Mini Kit | Qiagen | Cat#74104 |
| **Deposited data** | | |
| RNA-seq raw data and processed read counts | This paper | GEO: GSE313018 |
| ChIP-seq raw and processed data | This paper | GEO: GSE313100 |
| Microarray raw and processed data | This paper | GEO: GSE342162 |
| Global proteomic raw and processed data | This paper | PRIDE: PXD071760 |
| Phosphoproteomic raw and processed data | This paper | PRIDE: PXD071761 |
| Human reference genome NCBI build 38, GRCh38 | Genome Reference Consortium | http://www.ncbi.nlm.nih.gov/projects/genome/assembly/grc/human/ |
| Human cholangiocarcinoma single cell RNA data | Ma et al.[12]<br>Zhang et al.[13]<br>Massalha et al.[14]<br>Ma et al.[15] | GEO: GSE125449,<br>GEO: GSE138709,<br>GEO: GSE146409,<br>GEO: GSE151530. |
| Andersen cohort cholangiocarcinoma RNA data | Andersen et al.[5] | GEO: GSE26566 |
| Job cohort cholangiocarcinoma RNA data | Job et al.[8] | ArrayExpress: E-MTAB-6389 |
| TIGER cohort cholangiocarcinoma RNA data | Chaisaingmongkol et al.[10] | GEO: GSE76297 |
| Montal cohort cholangiocarcinoma RNA data | Montal et al.[11] | GEO: GSE132305 |
| Dong cohort cholangiocarcinoma RNA and proteomic data | Dong et al.[7] | NODE database: OEP001105 |
| Nakamura cohort cholangiocarcinoma RNA data | Nakamura et al.[28] | EGA: EGA00001000950 |
| pan-cancer transcriptomics | | TCGA datasets |
| CHOL cohorts in pan-cancer analysis | Job et al.[8]<br>Chaisaingmongkol et al.[10]<br>Nakamura et al.[28]<br>Montal et al.[11] | ArrayExpress: E-MTAB-6389<br>GEO: GSE76297<br>EGA:EGA00001000950<br>GEO: GSE132305 |
| Expression of in-house NHC and CCA cell lines | This paper | N/A |
| CRISPR-screen and copy number data in CCA cell lines | DepMap | https://depmap.org/portal/ |

| | | |
|---|---|---|
| Human cholangiocarcinoma spatial transcriptomic data | Gehl et al.[6] | N/A |
| **Experimental models: Cell lines** | | |
| Human: HUCCT-1 | Riken BRC Cell Bank | Cat# RCB1960 |
| Human: EGI-1 | DSMZ | Cat# ACC 385 |
| Human: SNU478 | KCLB | Cat# 00478 |
| Human: SNU1079 | KCLB | Cat# 01079 |
| Human: SNU1196 | KCLB | Cat# 01196 |
| Human: REB | Riken BRC Cell Bank | N/A |
| Human: WITT | Mayo Clinic, Rochester, MN | N/A |
| Human: KMCH | Mayo Clinic, Rochester, MN | N/A |
| Human: TKKK | RIKEN | N/A |
| Human: Huh28 | JCRB | N/A |
| Human: SNU308 | KCLB | Cat# 00308 |
| Human: CCLP1 | Donated by Dr. Ben J Dwyer | N/A |
| Human: SNU245 | KCLB | Cat# 00245 |
| Human: YSCCC | RIKEN | N/A |
| Human: SSP25 | RIKEN | N/A |
| Human: CCSW1 | Donated by Dr. Ben J Dwyer | N/A |
| Human: CC16 | Jens U. Marquardt, Universitatsklinikum Schleswig-Holstein (UKSH), Germany | PMID: 30485423 |
| Human: KMBC | Mayo Clinic | N/A |
| Human: immobilized cholangiocyte H69 | Mayo Clinic, Rochester, MN | N/A |
| Human: Primary cholangiocyte NHC2 | Jesus M. Banales, BioDonostia, Spain | PMID: 28237397 |
| Human: Primary cholangiocyte NHC3 | Jesus M. Banales, BioDonostia, Spain | PMID: 28237397 |
| Human: Primary cholangiocyte NHCSS | Jesus M. Banales, BioDonostia, Spain | PMID: 28237397 |
| Human: Primary cholangiocyte C324 | Jesus M. Banales, BioDonostia, Spain | PMID: 28237397 |
| **Experimental models: Organisms/strains** | | |
| Athymic NCr-nu/nu | Charles River Laboratories | Strain: #490 |
| **Oligonucleotides** | | |
| sgRNA oligos | IDT technologies | Table S8A |
| Primers used in this study | TAG Copenhagen A/S | Table S8B |
| **Recombinant DNA** | | |
| psPAX2 | Addgene | Cat#12260 |
| pCMV-VSV-G | Addgene | Cat#8454 |
| pSpCas9(BB)-2A-GFP (PX458) backbone | Addgene | Cat#48138 |
| pSpCas9-KDM3A-GFP | This paper | N/A |
| pSpCas9-KDM3B-GFP | This paper | N/A |
| pSpCas9-KDM5A-GFP | This paper | N/A |

| | | |
|---|---|---|
| pSpCas9-KDM5B-GFP | This paper | N/A |
| **Software and algorithms** | | |
| GraphPad Prism 9.3.1 | Graphpad | https://www.graphpad.com/ |
| R v4.5.1 | R Foundation | https://cran.rstudio.com/ |
| ImageJ | ImageJ | https://imagej.net/ij/ |
| Image Lab | Bio-Rad | https://www.bio-rad.com/en-dk/product/image-lab-software?ID=KRE6P5E8Z |
| BioRender | BioRender | https://www.biorender.com/ |
| Incucyte 2022B Rev1 | Sartorius | https://downloads.essenbioscience.com/get/incucyte-2022b-rev1-gui |
| DESeq2 v1.42.1 | Michael et al[29] | https://bioconductor.org/packages/release/bioc/html/DESeq2.html |
| nf-core/chipseq v2.0.0 | Ewels etal[18] | https://nf-co.re/chipseq/2.0.0/ |
| Bowtie2 v2.4.4 | Langmead et al[30] | https://bowtie-bio.sourceforge.net/bowtie2/index.shtml |
| Picard v2.27.4 | Broad Institute | https://github.com/broadinstitute/picard |
| MACS2 v2.2.7.1 | Zhang et al[31] | https://github.com/macs3-project/MACS/ |
| BEDTools | Quinlan et al[32] | https://bedtools.readthedocs.io/en/latest/ |
| Integrative Genomics Viewer (IGV) v2.15.4 | Robinson et al[33] | https://igv.org/ |
| deepTools v3.5.1 | Ramirez et al[34] | https://usegalaxy.eu/ |
| HOMER v4.11 | Heinz et al[35] | http://homer.ucsd.edu/homer/ |
| Spectronaut version 19.9.250422 (Biognosys) | Biognosys | https://biognosys.com/software/spectronaut/ |
| Clinical Knowledge Graph | Santos et al[19] | https://ckg.readthedocs.io/en/latest/ |
| PhosphoSitePlus | Hornbeck et al[36] | https://www.phosphosite.org/homeAction.action |
| Coral | Manning et al[37] | http://phanstiel-lab.med.unc.edu/CORAL/ |
| Seurat R package (v5.1.0) | Stuart et al[38] | https://cran.r-project.org/web/packages/Seurat/index.html |
| CellChat | Suoqin et al[26] | https://github.com/sqjin/CellChat |
| Cytoscape v3.10.2 | Cytoscape | https://cytoscape.org/download.html |

| | | |
|---|---|---|
| Gene Set Enrichment Analysis (GSEA) v4.3.3 | Broad Institute | https://www.gsea-msigdb.org/gsea/login.jsp |
| GSVA package v2.3.1 | Hänzelmann et al[39] | https://www.bioconductor.org/packages/release/bioc/html/GSVA.html |
| ggplot2 | Wickham et al[40] | https://cran.r-project.org/web/packages/ggplot2/index.html |
| Space Ranger v4.0.1 | 10x Genomics | https://www.10xgenomics.com/support/software/space-ranger/latest |
| ENACT v0.2.5 | Kamel et al[41] | https://github.com/Sanofi-Public/enact-pipeline |
| inferCNV v3.20 | Tirosh et al[42] | https://github.com/broadinstitute/infercnv |
| DBSCAN v1.7.2 | Schubert et al[43] | https://scikit-learn.org/stable/modules/generated/dbscan-function.html |
| SCOTIA v1.0 | Shiau et al[44] | https://github.com/Caochris/SCOTIA |
| igraph v2.0.3 | The igraph core team | https://github.com/igraph/rigraph |
| r-sf v1.0.14 | Pebesma et al[45] | https://r-spatial.github.io/sf |
| limma v2.10.7 | Ritchie et al[46] | https://github.com/gangwug/limma |
| Msigdbr v25.1.1 | Liberzon et al[47] | https://cran.r-project.org/web/packages/msigdbr/index.html |
| Other | | |

10.1038/srep05193.

of human intrahepatic cholangiocarcinoma. J Hepatol *73*, 1118-1130. 10.1016/j.jhep.2020.05.039.

14. Massalha, H., Halpern, K.B., Abu-Gazala, S., Jana, T., Massasa, E.E., Moor, A.E., Buchauer, L., Rozenberg, M., Pikarsky, E., Amit, I., et al. (2020). A single cell atlas of the human liver tumor microenvironment. Mol Syst Biol *16*. ARTN e9682

10.15252/msb.20209682.

15. Ma, L.C., Wang, L.M., Khatib, S.A., Chang, C.W., Heinrich, S., Dominguez, D.A., Forgues, M., Candia, J., Hernandez, M.O., Kelly, M., et al. (2021). Single-cell atlas of tumor cell evolution in response to therapy in hepatocellular carcinoma and intrahepatic cholangiocarcinoma. J Hepatol *75*, 1397-1408. 10.1016/j.jhep.2021.06.028.

16. Ramachandran, P., Dobie, R., Wilson-Kanamori, J.R., Dora, E.F., Henderson, B.E.P., Luu, N.T., Portman, J.R., Matchett, K.P., Brice, M., Marwick, J.A., et al. (2019). Resolving the fibrotic niche of human liver cirrhosis at single-cell level. Nature *575*, 512-+. 10.1038/s41586-019-1631-3.

17. Chen, E.Y., Tan, C.M., Kou, Y., Duan, Q.N., Wang, Z.C., Meirelles, G.V., Clark, N.R., and Ma'ayan, A. (2013). Enrichr: interactive and collaborative HTML5 gene list enrichment analysis tool. Bmc Bioinformatics *14*. Artn 128

10.1186/1471-2105-14-128.

18. Ewels, P.A., Peltzer, A., Fillinger, S., Patel, H., Alneberg, J., Wilm, A., Garcia, M.U., Di Tommaso, P., and Nahnsen, S. (2020). The nf-core framework for community-curated bioinformatics pipelines. Nature Biotechnology *38*, 276-278. 10.1038/s41587-020-0439-x.

19. Santos, A., Colaço, A.R., Nielsen, A.B., Niu, L.L., Strauss, M., Geyer, P.E., Coscia, F., Albrechtsen, N.J.W., Mundt, F., Jensen, L.J., and Mann, M. (2022). A knowledge graph to interpret clinical proteomics data. Nature Biotechnology *40*, 692-+. 10.1038/s41587-021-01145-6.

20. Lazar, C., Gatto, L., Ferro, M., Bruley, C., and Burger, T. (2016). Accounting for the Multiple Natures of Missing Values in Label-Free Quantitative Proteomics Data Sets to Compare Imputation Strategies. J Proteome Res *15*, 1116-1125. 10.1021/acs.jproteome.5b00981.

21. Hornbeck, P.V., Zhang, B., Murray, B., Kornhauser, J.M., Latham, V., and Skrzypek, E. (2015). PhosphoSitePlus, 2014: mutations, PTMs and recalibrations. Nucleic Acids Research *43*, D512-D520. 10.1093/nar/gku1267.

22. Gnad, F., Gunawardena, J., and Mann, M. (2011). PHOSIDA 2011: the posttranslational modification database. Nucleic Acids Research *39*, D253-D260. 10.1093/nar/gkq1159.

23. Johnson, J.L., Yaron, T.M., Huntsman, E.M., Kerelsky, A., Song, J.H., Regev, A., Lin, T.Y., Liberatore, K., Cizin, D.M., Cohen, B.M., et al. (2023). An atlas of substrate specificities for the human serine/threonine kinome. Nature *613*, 759-+. 10.1038/s41586-022-05575-3.

24. Yaron-Barir, T.M., Joughin, B.A., Huntsman, E.M., Kerelsky, A., Cizin, D.M., Cohen, B.M., Regev, A., Song, J., Vasan, N., Lin, T.Y., et al. (2024). The intrinsic substrate specificity of the human tyrosine kinome. Nature *629*, 1174-+. 10.1038/s41586-024-07407-y.

25. Metz, K.S., Deoudes, E.M., Berginski, M.E., Jimenez-Ruiz, I., Aksoy, B.A., Hammerbacher, J., Gomez, S.M., and Phanstiel, D.H. (2018). Coral: Clear and Customizable Visualization of Human Kinome Data. Cell Syst *7*, 347-+. 10.1016/j.cels.2018.07.001.

26. Jin, S.Q., Guerrero-Juarez, C.F., Zhang, L.H., Chang, I., Ramos, R., Kuan, C.H., Myung, P., Plikus, M.V., and Nie, Q. (2021). Inference and analysis of cell-cell communication using CellChat. Nature Communications *12*. ARTN 1088

10.1038/s41467-021-21246-9.

27. Xue, R.D., Zhang, Q.M., Cao, Q., Kong, R.R., Xiang, X., Liu, H.K., Feng, M., Wang, F.Y.N., Cheng, J.H., Li, Z., et al. (2022). Liver tumour immune microenvironment subtypes and neutrophil heterogeneity. Nature *612*, 141-+. 10.1038/s41586-022-05400-x.

28. Nakamura, H., Arai, Y., Totoki, Y., Shirota, T., Elzawahry, A., Kato, M., Hama, N., Hosoda, F., Urushidate, T., Ohashi, S., et al. (2015). Genomic spectra of biliary tract cancer. Nat Genet *47*, 1003-1010. 10.1038/ng.3375.

29. Love, M.I., Huber, W., and Anders, S. (2014). Moderated estimation of fold change and dispersion for RNA-seq data with DESeq2. Genome Biol *15*, 550. 10.1186/s13059-014-0550-8.

30. Langmead, B., and Salzberg, S.L. (2012). Fast gapped-read alignment with Bowtie 2. Nat Methods *9*, 357-U354. 10.1038/Nmeth.1923.

31. Zhang, Y., Liu, T., Meyer, C.A., Eeckhoute, J., Johnson, D.S., Bernstein, B.E., Nussbaum, C., Myers, R.M., Brown, M., Li, W., and Liu, X.S. (2008). Model-based Analysis of ChIP-Seq (MACS). Genome Biology *9*. ARTN R137

10.1186/gb-2008-9-9-r137.

32. Quinlan, A.R. (2014). BEDTools: The Swiss-Army Tool for Genome Feature Analysis. Curr Protoc Bioinformatics *47*, 11 12 11-34. 10.1002/0471250953.bi1112s47.

33. Robinson, J.T., Thorvaldsdottir, H., Turner, D., and Mesirov, J.P. (2023). igv.js: an embeddable JavaScript implementation of the Integrative Genomics Viewer (IGV). Bioinformatics *39*. ARTN btac830

10.1093/bioinformatics/btac830.

34. Ramirez, F., Ryan, D.P., Gruning, B., Bhardwaj, V., Kilpert, F., Richter, A.S., Heyne, S., Dundar, F., and Manke, T. (2016). deepTools2: a next generation web server for deep-sequencing data analysis. Nucleic Acids Res *44*, W160-165. 10.1093/nar/gkw257.

35. Heinz, S., Benner, C., Spann, N., Bertolino, E., Lin, Y.C., Laslo, P., Cheng, J.X., Murre, C., Singh, H., and Glass, C.K. (2010). Simple Combinations of Lineage-Determining Transcription Factors Prime

-Regulatory Elements Required for Macrophage and B Cell Identities. Molecular Cell *38*, 576-589. 10.1016/j.molcel.2010.05.004.

36. Hornbeck, P.V., Kornhauser, J.M., Tkachev, S., Zhang, B., Skrzypek, E., Murray, B., Latham, V., and Sullivan, M. (2012). PhosphoSitePlus: a comprehensive resource for investigating the structure and function of experimentally determined post-translational modifications in man and mouse. Nucleic Acids Research *40*, D261-D270. 10.1093/nar/gkr1122.

37. Manning, G., Whyte, D.B., Martinez, R., Hunter, T., and Sudarsanam, S. (2002). The protein kinase complement of the human genome. Science *298*, 1912-+. DOI 10.1126/science.1075762.

38. Stuart, T., Butler, A., Hoffman, P., Hafemeister, C., Papalexi, E., Mauck, W.M., Hao, Y.H., Stoeckius, M., Smibert, P., and Satija, R. (2019). Comprehensive Integration of Single-Cell Data. Cell *177*, 1888-+. 10.1016/j.cell.2019.05.031.

39. Hänzelmann, S., Castelo, R., and Guinney, J. (2013). GSVA: gene set variation analysis for microarray and RNA-Seq data. Bmc Bioinformatics *14*. Artn 7

10.1186/1471-2105-14-7.

40. Wickham, H. (2009). ggplot2: Elegant Graphics for Data Analysis. Use R, 1-212. 10.1007/978-0-387-98141-3.

41. Kamel, M., Song, Y., Solbas, A., Villordo, S., Sarangi, A., Senin, P., Sunaal, M., Ayestas, L.C., Levin, C., Wang, S.Q., et al. (2025). ENACT: End-to-End Analysis of Visium High Definition (HD) Data. Bioinformatics *41*. ARTN btaf094

10.1093/bioinformatics/btaf094.

42. Tirosh, I., Izar, B., Prakadan, S.M., Wadsworth, M., Treacy, D., Trombetta, J.J., Rotem, A., Rodman, C., Lian, C., Murphy, G., et al. (2016). Dissecting the multicellular ecosystem of

metastatic melanoma by single-cell RNA-seq. Science *352*, 189-196. 10.1126/science.aad0501.

43. Schubert, E., Sander, J., Ester, M., Kriegel, H.P., and Xu, X.W. (2017). DBSCAN Revisited, Revisited: Why and How You Should (Still) Use DBSCAN. Acm T Database Syst *42*. Artn 19

10.1145/3068335.

44. Cao, J.Y., Shiau, C., Gong, D., Gregory, M.T., Yin, X.Q., Cho, J.W., Wang, P.L., Su, J., Reeves, J.W., Guo, J.A., et al. (2024). Therapy-associated remodeling of pancreatic cancer revealed by single-cell spatial transcriptomics and optimal transport analysis. Cancer Research *84*. 10.1158/1538-7445.Am2024-3878.

45. Pebesma, E. (2018). Simple Features for R: Standardized Support for Spatial Vector Data. R J *10*, 439-446.

46. Ritchie, M.E., Phipson, B., Wu, D., Hu, Y.F., Law, C.W., Shi, W., and Smyth, G.K. (2015). Limma powers differential expression analyses for RNA-sequencing and microarray studies. Nucleic Acids Research *43*. ARTN e47

10.1093/nar/gkv007.

47. Liberzon, A., Birger, C., Thorvaldsdottir, H., Ghandi, M., Mesirov, J.P., and Tamayo, P. (2015). The Molecular Signatures Database Hallmark Gene Set Collection. Cell Syst *1*, 417-425. 10.1016/j.cels.2015.12.004.
